# Astrocyte CCN1 stabilizes neural circuits in the adult brain

**DOI:** 10.1101/2024.03.14.585077

**Authors:** Laura Sancho, Matthew M. Boisvert, Trinity Dawoodtabar, Jillybeth Burgado, Ellen Wang, Nicola J. Allen

## Abstract

Neural circuits in many brain regions are refined by experience. Sensory circuits support higher plasticity at younger ages during critical periods - times of circuit refinement and maturation - and limit plasticity in adulthood for circuit stability. The mechanisms underlying these differing plasticity levels and how they serve to maintain and stabilize the properties of sensory circuits remain largely unclear. By combining a transcriptomic approach with *ex vivo* electrophysiology and *in vivo* imaging techniques, we identify that astrocytes release cellular communication network factor 1 (CCN1) to maintain synapse and circuit stability in the visual cortex. By overexpressing CCN1 in critical period astrocytes, we find that it promotes the maturation of inhibitory circuits and limits ocular dominance plasticity. Conversely, by knocking out astrocyte CCN1 in adults, binocular circuits are destabilized. These studies establish CCN1 as a novel astrocyte-secreted factor that stabilizes neuronal circuits. Moreover, they demonstrate that the composition and properties of sensory circuits require ongoing maintenance in adulthood, and that these maintenance cues are provided by astrocytes.

## Main Text

Mammalian neural circuits are highly adaptable at an early age as the organism learns to adapt to its environment. This is evident in sensory circuits, which typically have higher neural plasticity at younger ages, a period of circuit refinement, and increased stability and reduced plasticity in adulthood (*1, 2*). This stability in adulthood maintains neuronal circuit connections (*3*). However, the extent to which sensory circuit properties and synapses are actively stabilized in adulthood remains unclear. Astrocytes are ideal candidates for dynamically regulating sensory circuit plasticity and stability in adulthood. Astrocytes are a major glial subtype that in adulthood support neuronal function through multiple mechanisms, including regulating ion homeostasis, neuronal metabolism, nutrient supply, and neurotransmitter reuptake (*4–9*). While previous studies have implicated astrocytes and astrocyte-secreted factors in regulating synapse formation, function, and stabilization (*10–12*), the mechanisms by which astrocytes actively maintain sensory circuit stability and limit plasticity in the adult brain remain unclear.

The mouse visual cortex is an ideal circuit for examining how multiple components of circuit and synapse stability are maintained by astrocytes. It has well-characterized stereotyped periods of high plasticity early in development and high stability in adulthood (1, 2). During the critical period for binocular vision, which in mice begins at approximately the end of the third postnatal week and lasts for two weeks, essential properties of the visual system are established. These include ocular dominance, the preference of a neuron to input from one eye compared to the other (*13–16*). By the end of the critical period the binocular circuitry is well-established and fine-tuned (3). However, this stability in adulthood can be reversible, as digesting the extracellular matrix with enzymes or transplanting juvenile inhibitory interneurons can re-open the window for ocular dominance plasticity (*17, 18*).

The developmental maturation of astrocytes in the visual cortex follows that of the neurons they interact with, with astrocytes producing stage-specific cues that instruct synapse formation, maturation, and elimination (*9, 19*). Thus, astrocytes are well positioned to regulate plasticity state and hence synaptic stability. Early experiments showed that transplantation of juvenile astrocytes into adult visual cortex can reopen the window for plasticity (*12*). Astrocytes can also impact plasticity through chordin-like 1 and connexin 30 (*10, 11*). However, the mechanisms by which astrocytes close the window for plasticity and promote stability in the adult brain are largely unexplored.

To identify astrocyte factors underlying periods of high plasticity vs high synapse stability, we conducted a transcriptomic screen in visual cortex astrocytes across key developmental timepoints and after plasticity-inducing manipulations. We establish a novel astrocyte secreted factor, cellular communication network factor 1 (CCN1), as a factor that regulates circuit stability in mouse visual cortex. CCN1 is a 4-domain secreted protein that can bind with many components of the plasma membrane and the extracellular matrix (ECM), including heparan sulfate proteoglycans, integrins, and connexins (*20, 21*). The role of CCN1 in the periphery has been investigated in the context of tumorigenesis, angiogenesis, inflammation, and lung development (*22–27*), though its role in the central nervous system (CNS) remains largely unknown. In this study, we show that astrocytic CCN1 regulates circuit stability by exerting its effect on multiple cell types, including excitatory neurons, inhibitory neurons, and microglia.

### Transcriptional programs in astrocytes underlying synapse and circuit stability

To determine whether astrocytes change their synapse- and circuit-regulating properties across periods of high synaptic plasticity and high synaptic stability, we performed bulk RNA sequencing of visual cortex astrocytes using astrocyte Ribotag (Rpl22^fl/fl^;GFAPcre) mice (Fig. 1A, Supplemental Data Table 1) (*19, 28*). This experiment was designed to identify “pro-stability” genes, those that are upregulated in astrocytes in adulthood (P120) compared to early development and the critical period (P28), and downregulated after plasticity-inducing paradigms. We performed transcriptional profiling of visual cortex astrocytes at different developmental stages and after plasticity-inducing paradigms: 1) dark rearing until P45 and 2) two days of monocular deprivation (MD) at P28 (Fig. 1B) (*19*). Dark rearing delays astrocyte maturation and the closure of the visual critical period (*29–31*) while MD during the critical period induces ocular dominance plasticity (*14*). Critical period (P28) and adult (P120) data was obtained from a previously published study (*19, 28*).

**Figure 1.**
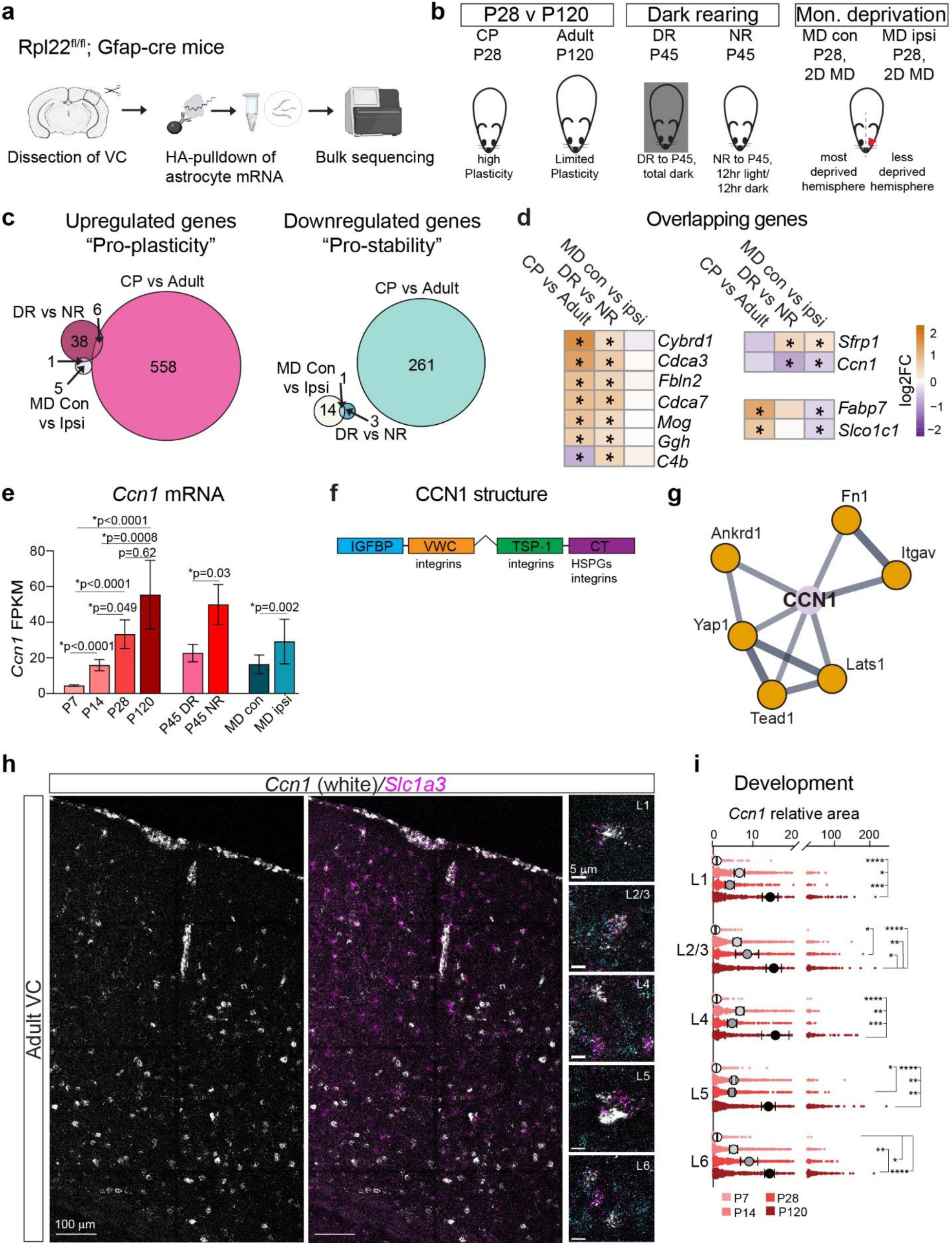
Transcriptional programs in astrocytes underlying synapse and circuit stability. **a**. Schematic of experimental setup for bulk transcriptomics. Visual cortices (VC) of Rpl22fl/fl x GFAP-cre mice were dissected, an HA-pulldown of astrocyte mRNA was performed, and bulk sequencing was run. **b**. Schematic of different experimental conditions. CP, critical period. MD, monocular deprivation (lid suturing). Con, contralateral eye to MD. Ipsi, ipsilateral eye to MD. DR, dark rearing. NR, normal rearing. **c**. Venn diagram of overlapping astrocyte differentially expressed genes (DEGs) in different experimental comparisons. Criteria: Fragments per kilobase of transcript per million mapped reads (FPKM) > 1 in at least one group per comparison, ribotag pulldown FPKM (astrocyte) /input FPKM (all cells) > 0.75 in each at least one group per comparison, adjusted p-value < 0.05, Log2 Fold Change (LFC) > |0|). **d**. Heatmap of the DEGs in more than one comparison from **c**. Adjusted p-value < 0.05 in at least 2 comparisons. Color scale is -2.3 to 2.3 LFC. Stars denote significant comparison. **e**. FPKM for *Ccn1* in different experimental conditions. Developmental data from Farhy-Tselnicker et al., 2021 and Boisvert et al., 2018. Statistics done with DESeq2, error bars show SEM. **f**. Protein structure of CCN1 with 4 domains showing binding sites for integrins and heparan sulfate proteoglycans (HSPGs). **g**. Predicted functional protein network of CCN1 using STRING database. **h**. Representative images of P120 smFISH in a tile image of the VC (left) and in different cortical layers within the VC (right). smFISH against *Slc1a3* and *Ccn1.* Scale bar, 5 µm. **i**. *Ccn1* thresholded area within *Slc1a3* regions of interest (astrocytes) in different cortical layers in the VC at P7, P14, P28, P120. Small dots denote individual astrocytes, large circles are averages by mouse with error bars showing SEM. N = 3 mice per age. 2-way ANOVA with post-hoc Tukey tests.

We determined differentially expressed genes (DEGs) that were enriched in astrocytes in three comparisons - dark rearing vs normal rearing, MD contralateral to the deprived eye vs MD ipsilateral, and P28 (critical period) vs P120 (adult) (Fig. 1B and C, Extended Data Fig.1A). Gene-set enrichment analysis (GSEA) using the Reactome database identified the involvement of DEGs in lipid and steroid metabolic processes and extracellular matrix organization, generally with upregulation in higher plasticity conditions such as during the critical period and after deprivation in the contralateral hemisphere (Extended Data Fig.1B; Supplemental Data Table 2). Overrepresentation analysis (ORA) revealed the role of DEGs in synapse structure, membrane organization, and cell adhesion (Extended Data Fig.2, Extended Data Fig.3A; Supplemental Data Table 3). To identify pro-stability candidate genes, we focused on genes altered in more than one comparison (Fig. 1D; 11 genes). We hypothesized that an astrocytic pro-stability gene would be upregulated throughout development, stay high in adulthood, and be downregulated after dark rearing and MD. A promising candidate that fits this profile is cellular communication network factor 1 (*Ccn1*) also known as cysteine-rich angiogenic inducer 61 (*Cyr61*) (Fig. 1E). *Ccn1* encodes a four-domain secreted protein, CCN1, that contains integrin and heparan sulfate proteoglycan (HSPGs) binding sites (Fig. 1F) and whose predicted functional interaction network highlights its association with integrins (*Itgav*), the extracellular matrix, and different transcriptional regulators (Fig. 1G), highlighting its potential role as a regulator of neuronal plasticity and stability.

To validate the expression profile of *Ccn1* detected by RNA sequencing, we used single molecule fluorescent in situ hybridization (smFISH) to characterize *Ccn1* in visual cortex astrocytes across different ages and plasticity paradigms. This shows *Ccn1* is expressed by astrocytes in all layers of the visual cortex, and expression increases across development and stays high into adulthood (Fig. 1H and I, Extended Data Fig.3B). *Ccn1* is also expressed by neurons, though at lower level (*32*) (Extended Data Fig.3C). We validated that dark-rearing decreases *Ccn1* in astrocytes (Extended Data Fig.3D). Thus, the expression pattern of *Ccn1*, being high in adulthood and low after plasticity paradigms, suggests that CCN1 may be acting as a “pro-stability” factor produced by astrocytes.

### Astrocyte CCN1 restricts large-scale binocular zone remodeling

To test the role of CCN1 as a pro-stability factor, we manipulated astrocyte expression of *Ccn1 in vivo*. We overexpressed CCN1 in astrocytes of the visual cortex using viral vectors at a time when plasticity is high, the juvenile critical period, to ask if we could prematurely restrict plasticity. We injected the visual cortex of wild-type (WT) mice with AAV2/5-tdTomato as a control (tdT) or AAV2/5-CCN1-HA (CCN1) driven by the minimal glial fibrillary acidic protein (GFAP) promoter at P14 (Fig. 2A). We validated that the GFAP-CCN1-HA plasmid led to expression of HA-tagged CCN1 protein using cultured astrocytes and HEK cells (*33*) (Extended Data Fig.4A and B). These constructs were specific for astrocytes and had high penetrance (Fig. 2B and C) and did not induce differential glial reactivity (Extended Data Fig.4C and D). To probe for plasticity, we performed monocular enucleation (ME) to induce remodeling of inputs to the visual cortex, with outcomes measured with an *Arc* induction assay (*34, 35*) (Fig. 2D). The width of *Arc* expression, an immediate early gene expressed in visual cortex neurons activated by light, can approximate the size of the binocular zone (BZ) (*10, 16*). Four days of monocular enucleation in mice during the critical period typically results in an expansion of the *Arc* activation width in the hemisphere ipsilateral to the open eye relative to control overnight enucleated mice (*16*) (Fig. 2D). When we performed these experiments at P28, the peak of the critical period, in mice overexpressing (o/e) tdT or CCN1 in astrocytes, we found that CCN1 o/e mice had less binocular zone remodeling after four days of monocular enucleation, relative to tdT o/e mice, as measured by width of the *Arc* zone (Fig. 2E and F, Extended Data Fig.5A). This finding indicates that overexpressing CCN1 actively restricts plasticity at a time when it is normally high, thus promoting binocular zone stability.

**Figure 2.**
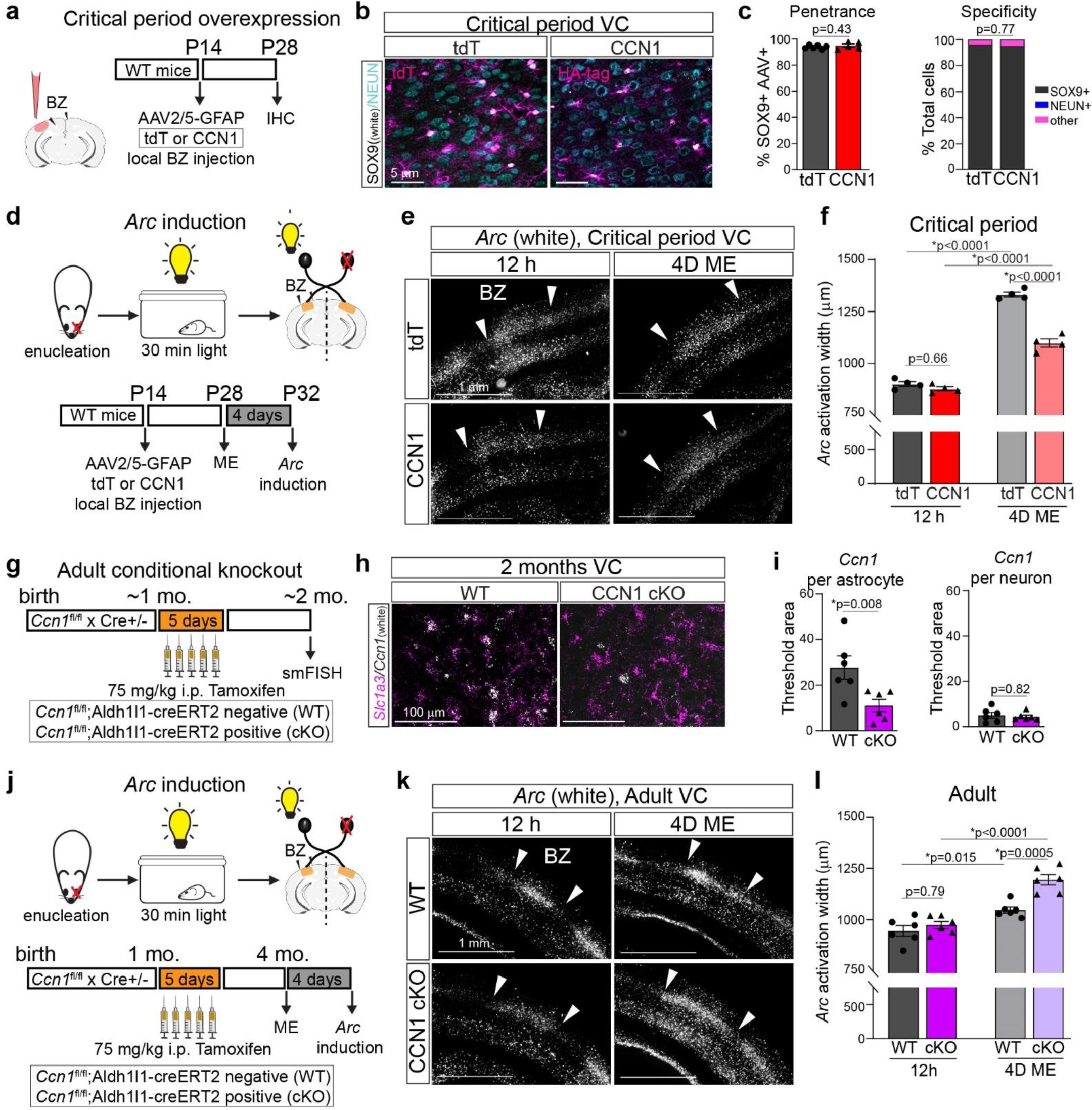
Astrocytic CCN1 restricts large-scale binocular zone remodeling. **a**. Schematic of experimental set-up for critical period overexpression of CCN1 or tdTomato (tdT) viral vectors in astrocytes. BZ, binocular zone. **b**. Representative images of P28 VC expressing tdT (left) or CCN1 (right). Scale bar, 50 µm. **c**. Left, penetrance of viral vectors quantified as % SOX9+ cells also expressing either tdT or HA-tag for CCN1 group. Right, specificity quantified as % total cells expressing tdT or CCN1 that are co-stained with SOX9, NEUN or no staining (other). tdT, 6 mice. CCN1, 5 mice. Unpaired T-test. **d**. Top, schematic of *Arc* induction assay. Bottom, schematic of experimental timeline for critical period mice. **e**. Representative image of *Arc* smFISH in the BZ. Left, 12 h after ME, monocular enucleation. Right, 4 days after ME (4D ME). Scale bar, 1 mm. White arrows denote *Arc* BZ width. **f**. Quantification of *Arc* activation width in layer 4 of hemisphere contralateral to enucleated eye. N = 4 mice per group. Two-way ANOVA with post-hoc Tukey tests. **g**. Schematic of experimental set-up for adult cKO of CCN1 in astrocytes. **h**. Representative image of 2 month VC smFISH against *Slc1a3* and *Ccn1*. Scale bar, 100 µm. **i**. Left, quantification of *Ccn1* threshold area per astrocyte. Unpaired T-test. Right, *Ccn1* threshold area per neuron (labeled with *Tubb3*, bottom). Mann-Whitney U-test. N = 6 mice per genotype. **j**. Top, schematic of *Arc* induction assay. Bottom, schematic of experimental timeline in adult CCN1 cKO mice. **k**. Representative images as in **e**. Scale bar, 1 mm. **l**. Quantification of *Arc* width as in **f**. N = 6 mice per group. Two-way ANOVA with post-hoc Tukey tests. Bar charts show mean ± SEM, data points mice.

Conversely, we hypothesized that removing CCN1 from astrocytes in adulthood would result in increased plasticity. To knock out CCN1 specifically in astrocytes, we crossed a *Ccn1* floxed mouse (*36*) with a tamoxifen-inducible Aldh1l1-CreERT2 mouse (*37*), with tamoxifen given after 1 month of age (Fig. 2G). smFISH (Fig. 2H) demonstrated that the cKO (*Ccn1*^fl/fl^; Cre+) is specific to astrocytes and reduces *Ccn1* mRNA expression in astrocytes by ∼60% on average relative to WT mice (*Ccn1*^fl/fl^; Cre-) with no change in neuronal expression (Fig. 2I and Extended Data Fig.4E and F). We performed monocular enucleation in P120 WT and CCN1 cKO mice, reflecting an adult timepoint when plasticity is low, and assessed the outcome using the *Arc* induction assay (Fig. 2J). This revealed increased plasticity after four days of monocular enucleation in CCN1 cKO mice (Fig. 2K and L, Extended Data Fig.5B), demonstrating that CCN1 is actively promoting visual cortex stability in adulthood. Overall, these findings identify a role for astrocyte-secreted CCN1 in restricting synaptic plasticity in visual circuits.

### Astrocyte CCN1 regulates synaptic input onto excitatory and inhibitory neurons in the visual cortex

To understand if CCN1 is impacting circuit stability by altering synapses, we performed *ex vivo* whole-cell patch clamp recordings of spontaneous excitatory postsynaptic currents (sEPSCs) in layer 2/3 pyramidal neurons and fast-spiking neurons (putative parvalbumin positive, PV, interneurons) (Fig. 3A and B). These recordings were performed in critical period age mice at P33-34 that were overexpressing tdT or CCN1, and underwent 5 days of MD (5D MD) or no manipulation (Fig. 3A). When we recorded from pyramidal neurons (Fig. 3C), we found an increase in the sEPSC interevent interval (i.e., a decrease in frequency) in CCN1 o/e mice relative to tdT o/e mice at baseline, which was reversed with MD (Fig. 3D and Extended Data Fig.6A). We found a significant main effect of viral group on sEPSC amplitude, with CCN1 o/e mice having smaller sEPSC amplitude relative to tdT o/e mice (Fig. 3E and Extended Data Fig.6A), with no change in sEPSC area (Fig. 3F), and a slower sEPSC decay in CCN1 o/e mice (Extended Data Fig.6B), regardless of manipulation. Overall, these data suggest that CCN1 decreases excitatory drive onto pyramidal neurons and that MD reverses some of these changes.

**Figure 3.**
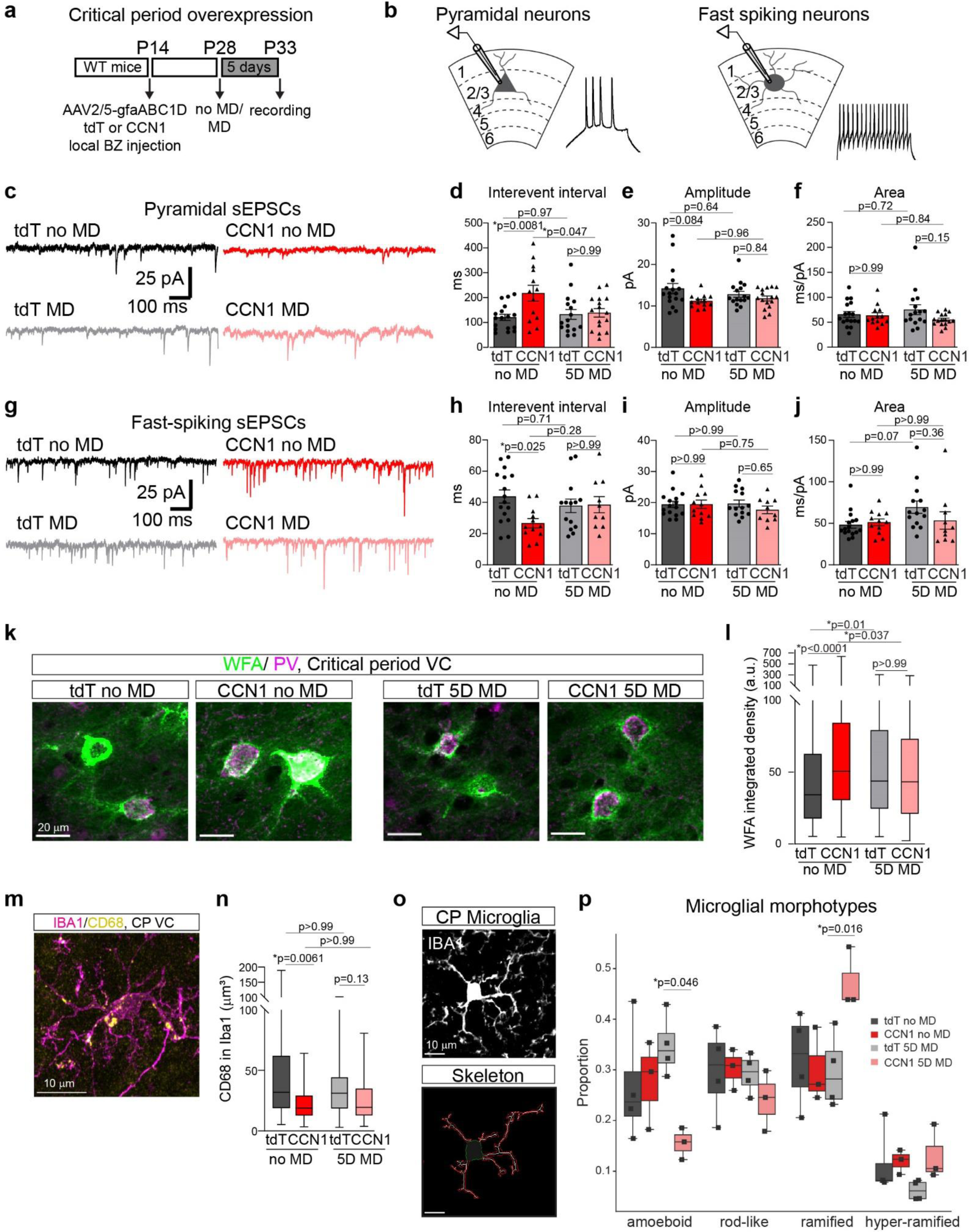
Astrocytic CCN1 regulates synaptic input on excitatory and inhibitory neurons in the visual cortex. **a**. Schematic of experimental timeline of viral vector injections and electrophysiology experiments in critical period mice. **b**. Whole-cell patch clamp recordings of layer 2/3 pyramidal neurons (left) or fast spiking neurons (right). **c**. Representative spontaneous excitatory postsynaptic currents (sEPSCs) from each of the four conditions. **d**. Average interevent interval (ms). **e**. Average amplitude (pA). **f**. Average area (ms/pA). **d-f**. Black symbols denote individual cell averages. tdT no MD: n = 17 cells, 9 mice. tdT MD: n = 16 cells, 7 mice. CCN1 no MD: n = 13 cells, 7 mice. CCN1 MD: n = 16 cells, 6 mice. **g**. Representative sEPSCs as in **c** from fast-spiking neurons. **h**. Average interevent interval (ms). **i**. Average amplitude (pA). **j**. Average area (ms/pA). **h-j**. Black symbols denote individual cell averages. tdT no MD: n = 16 cells, 10 mice. tdT MD: n = 14 cells, 9 mice. CCN1 no MD: n = 12 cells, 7 mice. CCN1 MD: n = 10 cells, 6 mice. Two-way ANOVA with post-hoc Tukey tests. **k**. Representative images of wisteria floribunda agglutinin (WFA) and parvalbumin (PV) staining in critical period VC. Scale bar, 20 µm. **l**. WFA integrated density around PV+ cells. 5 mice per group. Tdt no MD, n = 204 cells. tdT MD, n = 197 cells. CCN1 no MD, n = 228 cells. CCN1 MD, n = 193 cells. Kruskal Wallis test with post-hoc Dunn’s tests. **m.** Representative image of IBA1and CD68 staining in critical period VC. Scale bar, 10 µm**. n.** Volume of CD68+ puncta within IBA1 volume. Kruskal Wallis test with post-hoc Dunn’s tests. **o**. Representative image of IBA1 staining from critical period VC, top. Representative cell body and process tracing (skeleton), bottom. Scale bar, 10 µm. tdT no MD, 747 microglia from 4 mice. tdT MD, 876 microglia from 4 mice. CCN1 no MD, 496 microglia from 3 mice. CCN1 MD, 379 microglia from 3 mice. **p**. Proportion of microglial morphotypes-amoeboid, rod-like, ramified, and hyper-ramified. Black symbols, mouse averages. Two-way ANOVA with post-hoc Tukey tests. Only significant tests are shown. Bar charts show mean ± SEM. Box charts show median, upper and lower quartiles and whiskers show full range of data.

Recording sEPSCs from fast-spiking neurons (Fig. 3G), which we identified morphologically and electrophysiologically (*38*), revealed that CCN1 o/e had the opposite effects to those in pyramidal neurons. While CCN1 o/e had no impact on sEPSC amplitude, we found a significant decrease in interevent interval, i.e. an increase in sEPSC frequency, in CCN1 o/e mice relative to tdT mice at baseline, indicating an increase in excitatory input (Fig. 3H and I; Extended Data Fig. 6F). We found no change in sEPSC area across groups (Fig. 3J), though there was a significant increase in the decay and risetime of sEPSCs in tdT o/e mice after MD, but not CCN1 o/e (Extended Data Fig.6G and H). This demonstrates that the overall impact of CCN1 o/e on synaptic networks is to decrease excitatory drive onto excitatory neurons, while increasing excitatory drive onto inhibitory neurons. This has important implications for circuit function as the increase of inhibitory circuit activity is thought to underlie the closure of the critical period and the stabilization of visual circuits (*1*).

### Astrocyte CCN1 promotes inhibitory circuit maturity and alters microglial state

A primary mechanism underlying the loss of plasticity in the visual cortex is the maturation of perineuronal nets (PNNs), which are dense extracellular matrices that form predominantly around parvalbumin (PV) interneurons (*2, 39–43*). As we found a decrease in binocular zone remodeling in CCN1 overexpressing critical period mice (Fig. 2E and F) and an increase in excitatory drive onto fast-spiking neurons (putative PV neurons; Fig. 3H), we hypothesized that critical period CCN1 o/e mice had premature maturation of PNNs. To test this, we performed immunohistochemistry against PV and labeled PNNs using wisteria floribunda agglutinin (WFA) in the visual cortex of tdT and CCN1 o/e mice at baseline and after 5 days of MD. At baseline CCN1 o/e mice had a pronounced increase in WFA signal around PV neurons, and around all cells, compared to tdT o/e mice across the visual cortex (Fig. 3K and L, Extended Data Fig.7A). In CCN o/e mice WFA was reduced after MD, while in tdT o/e mice there was an increase in PNN density around PV interneurons after MD (Fig. 3L). The extracellular matrix and PNNs can be remodeled by microglia in the CNS in response to plasticity and disease (*44–46*); additionally, CCN1 secretion activates macrophages in the periphery (*22*) while loss of CCN1 increases microglial activation in the retina (*47*). To assess whether these CCN1-dependent changes in PNN density are mediated by microglia, we performed super-resolution confocal imaging of microglia and PNNs. Microglia were labeled with IBA1, phagocytotic lysosomes were labeled with CD68, and PNNs were labeled with WFA (*46, 48*) (Extended Data Fig.7B). We found no difference in the colocalization of WFA and CD68 puncta contained within IBA1 positive microglia between tdT or CCN1 o/e mice before or after MD (Extended Data Fig.7C), suggesting that phagocytosis of PNNs by microglia does not play a role in this context.

To examine microglial activation state, we quantified CD68 volume within microglia and found that at baseline overexpressing CCN1 in astrocytes led to a decrease in CD68 volume within microglia relative to tdT o/e mice (Fig. 3M and N). We further characterized microglial state by tracing the processes and soma of IBA1-labeled microglia to create skeletonized microglia and performed detailed analysis of different morphological features (Methods; Fig. 3O). Dimensionality reduction via principal components analysis and k-means clustering were performed (Methods) to classify microglia into four morphotypes – amoeboid, rod-like, ramified, and hyper-ramified (*49–51*) (Extended Data Fig.8A and C). Microglial morphology is directly related to their function, with amoeboid and rod-like morphologies thought to reflect more phagocytotic states, ramified morphologies reflecting a homeostatic role, and hyper-ramified reflecting a response to stress or loss of sensory input (*52, 53*). We found that 5 days of MD led to no changes in the relative proportions of microglial morphotypes in tdT mice, where there was a lower proportion of hyper-ramified microglia at baseline relative to other morphotypes (Extended Data Fig.8B). CCN1 o/e mice at baseline also showed a lower proportion of the hyper-ramified morphotype. However, 5 days of MD in CCN1 o/e mice eliminated this difference and increased the relative proportion of ramified (resting) microglia (Extended Data Fig.8B). Additionally, MD in CCN1 o/e mice results in a decreased proportion of amoeboid microglia and an increase in ramified microglia relative to tdT mice after MD (Fig. 3P). These results indicate that MD induces changes to microglia morphology in CCN1 o/e mice that reflect a more ramified state even after 5 days of MD; this is also supported by an increase in the total skeleton length and mean branch length and decrease in form factor (Extended Data Fig.8D-G).

Taken together, these data demonstrate that CCN1 promotes the maturity of PNNs surrounding inhibitory interneurons, and that this increased PNN density at baseline is not likely mediated by reduced microglial engulfment of PNNs. We also found that CCN1 can regulate microglial state, as we observe a reduction in phagocytotic lysosomes and associated microglial morphotypes in CCN1 o/e mice, particularly after MD. This suggests an altered microglial state in the visual cortex with CCN1 overexpression that may have important implications for regulating plasticity and stability of synapses and circuits.

### Astrocyte CCN1 promotes binocular circuit composition and stability in adulthood *in vivo*

In adult mice, visual circuit properties are well characterized, yet the mechanisms regulating their stability remain unclear. To address the role of CCN1 in maintaining visual circuit functional stability, we used two-photon *in vivo* microscopy to examine binocular circuits in adult (P120) WT and CCN1 cKO mice. WT and cKO mice were implanted with a cranial window and injected with an AAV2/1 viral vector encoding a green fluorescent calcium indicator preferentially targeted to excitatory neurons (CAMKIIa-gCaMP6f) at three months of age (Fig. 4A). Mice were allowed to recover and habituated to head fixation on a linear treadmill (Fig. 4B). We imaged calcium responses in layer 2/3 pyramidal neurons in response to drifting sinusoidal gratings independently presented to each eye, and imaged regions were confirmed to be in the binocular zone by performing retinotopic mapping (*54*) (Fig. 4B, Extended Data Fig.9A-C; Methods).

**Figure 4.**
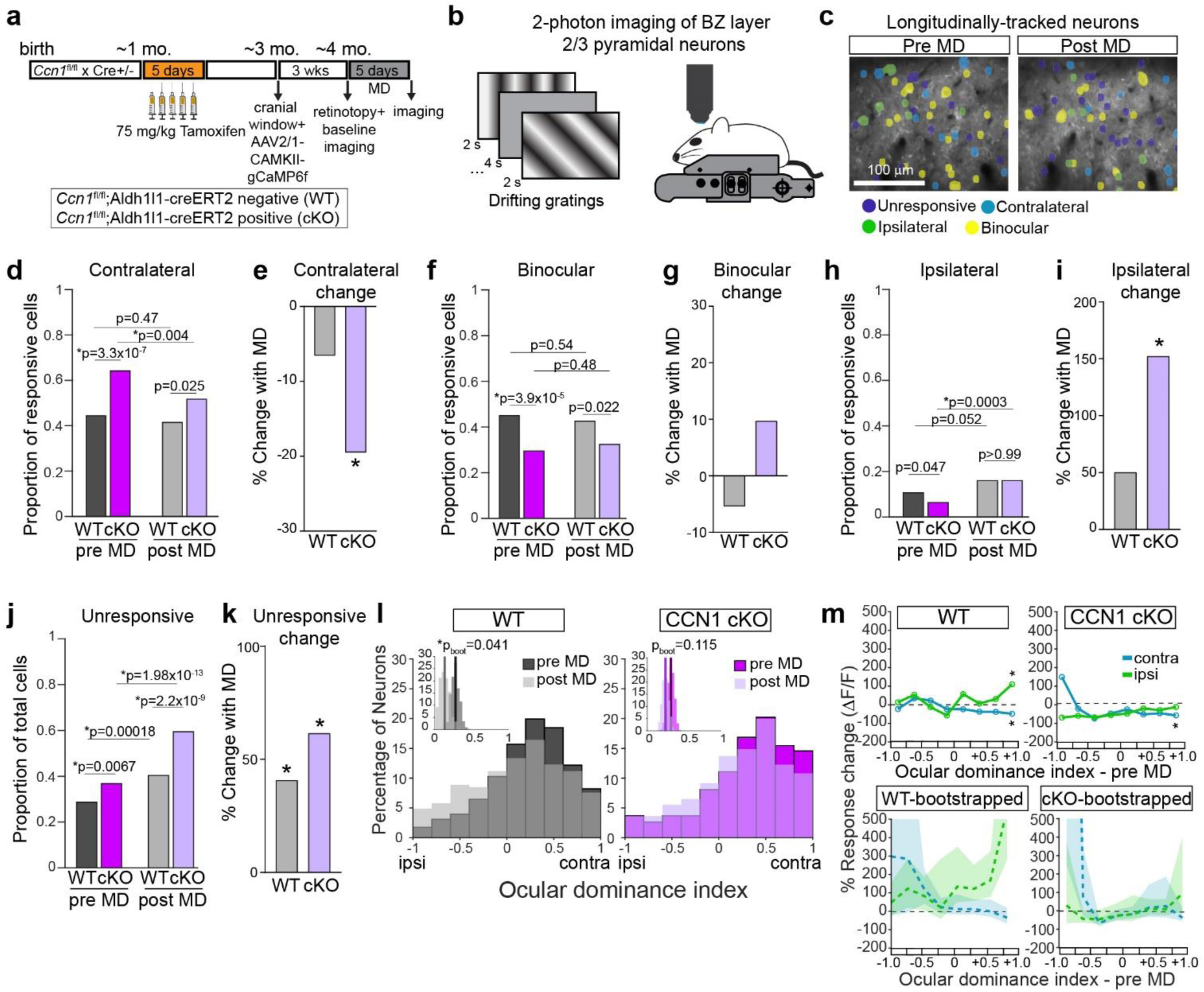
Astrocytic CCN1 promotes binocular circuit composition and stability in adulthood *in vivo*. **a**. Timeline of experiments. **b**. Schematic of 2-photon imaging of layer 2/3 pyramidal neurons in the binocular zone (BZ). Drifting sinusoidal gratings are presented independently to each eye in pseudorandom order to a head-fixed mouse for 2 s followed by 4 s of a gray screen. **c**. Representative image of longitudinal tracking of cell identity. Scale bar, 100 µm. **d**. Contralateral cells as a proportion of responsive cells. **e**. Change in contralateral cell proportion after MD, data and statistics from **d**. Star indicates significant change relative to pre MD. **f**. Binocular cells as a proportion of responsive cells. **g**. Change in binocular cell proportion after MD, data and statistics from **f**. Star indicates significant change relative to pre MD. **h**. Ipsilateral cells as a proportion of responsive cells. **i**. Change in ipsilateral cell proportion after MD, data and statistics from **h**. Star indicates significant change relative to pre MD. **d-i**. WT responsive cells pre MD = 329. WT responsive cells post MD = 275. CCN1 cKO responsive cells pre MD = 332. CCN1 cKO responsive cells post MD = 213. **j**. Unresponsive cells as a proportion of total cells. **k**. Change in unresponsive cell proportion after MD, data and statistics from **j**. Star indicates significant change relative to pre MD. **j-k.** WT total longitudinally tracked cells = 461. CCN1 cKO total longitudinally tracked cells = 525. **l**. Histogram of ocular dominance index (ODI) as a percentage of all responsive neurons, not necessarily longitudinally tracked. Inset is resampled data via hierarchical bootstrapping, showing significant leftward shift in ODI. All responsive cells: WT-pre MD = 682 cells, WT-post MD = 554 cells, cKO-pre MD = 569 cells, cKO-post MD = 736 cells. **m**. Longitudinally tracked responsive neurons only (responsive pre and post MD). WT = 223 cells. cKO = 163 cells. Percent magnitude response change in contralateral eye responses or ipsilateral eye responses. 8 bins based on pre MD ODI. Median contralateral or ipsilateral eye response changes per bin are denoted as circles. Stars indicated significant difference from 0% change, with multiple comparisons correction (new p-value threshold = 0.00625). Two-side Wilcoxon Rank Signed test. Top, experimental data. Bottom, bootstrapped means (dashed line) with a 95% confidence interval (shaded region). For **d-k**, chi-square test of proportions with correction for multiple comparisons (new p-value threshold = 0.0125, starred p-values denote significant values). Data from 4 mice for WT and 4 mice for cKO.

To characterize changes in binocular circuitry induced by monocular deprivation, we performed longitudinal tracking of layer 2/3 pyramidal neurons before and after 5 days of MD (Extended Data Fig.9D and F, Methods). Neurons were classified based on whether they were contralateral eye responsive, ipsilateral eye responsive, binocular, or unresponsive (*55–57*) (Fig. 4C; Methods). We found that CCN1 cKO mice had a higher proportion of contralateral responsive neurons relative to WT before MD, and that this proportion decreased with MD of the contralateral eye (Fig. 4D and E). These lost contralateral cells became ipsilateral, binocular, or unresponsive (Extended Data Fig.10A). We also found that CCN1 cKO mice had a smaller proportion of binocular neurons relative to WT before MD, and no significant change with MD (Fig. 4F and G) and an increased binocular cell turnover (Extended Data Fig.10B). Paralleling our results with the *Arc* induction assay (Fig. 2), which assesses neurons responsive to ipsilateral eye stimulation, we found that CCN1 cKO mice showed an increase in ipsilateral eye responsive neurons after MD, with no significant change in WT (Fig. 4H and I; Extended Data Fig.10C). After MD, both WT and CCN1 cKO showed an increase in unresponsive neurons, though this was markedly greater in cKO mice (Fig. 4J and K; Extended Data Fig.10D). This increase in unresponsiveness is similar to what is seen after monocular deprivation during the critical period in WT mice (*58*). Immature visual circuits have higher proportions of contralaterally responsive cells and lower proportions of binocularly responsive cells and these proportions change during the critical period (*55*), suggesting that astrocyte CCN1 is necessary for the maturity of binocular visual circuits.

To assess ocular dominance plasticity, ocular dominance mapping was performed before and after 5 days monocular deprivation of the contralateral eye. To determine whether MD had a differential effect on WT or CCN1 cKO ocular dominance within each cell, we calculated ocular dominance index (ODI) of all responsive neurons using the peak responses to the preferred stimuli for contralateral and ipsilateral eye stimulation (Fig. 4L; Extended Data Fig.10E-G; Methods). Due to the variability in ODI and the nested nature of the data, we performed hierarchical bootstrapping (*59*). We found a robust effect of MD in WT mice, with the mean ODI shifting towards the ipsilateral, non-deprived eye, but no change in CCN1 cKO mice (Fig. 4L). To verify this, we compared ODI from only the responsive, longitudinally tracked neurons and found a shift only in WT mice towards the ipsilateral eye (Extended Data Fig.10H). To determine whether ipsilateral or contralateral response magnitude changes were underlying the decrease in ODI, we plotted the changes in contralateral and ipsilateral eye responses for each longitudinally tracked responsive neuron as a function of its pre-MD ODI. We found that neurons with initially stronger contralateral eye biases experienced a significant decrease in contralateral eye response magnitudes in both WT and CCN1 cKO mice after MD (Fig. 4M). However, only WT mice experienced a concomitant increase in ipsilateral response magnitude. Bootstrapping verified the robustness of the ipsilateral response magnitude increase in WT neurons, with neurons with stronger contralateral eye biases before MD being more affected (Fig. 4M). Thus, monocular deprivation induces an increase in ipsilateral eye response magnitude in WT mice, but an increase in the proportion of ipsilateral responsive neurons in CCN1 cKO mice. These findings are unexpected and suggest that while WT mice undergo response magnitude changes to visual stimulation after MD, CCN1 cKO mice undergo a functional reorganization of their binocular circuitry after MD.

To establish whether alterations in the composition of the binocular circuitry that occurred in CCN1 cKO mice after 5 days of MD were due to the manipulation, or an intrinsic instability present in CCN1 cKO mice, we performed these ocular dominance and binocular circuitry composition measurements in naïve mice without MD at day 1 and day 6 of imaging. As before, we found that CCN1 cKO mice had more contralateral responsive cells and less binocular cells, with no significant changes between day 1 and day 6 of imaging, though with a trend for a decrease in binocular cells as has been previously reported in adult layer 2/3 pyramidal neurons over the course of one week of imaging (Extended Data Fig.11A and B) (*55*). We found no change in ipsilateral cells in the WT and cKO and no change between day 1 and day 6 of imaging (Extended Data Fig.11C). Because we saw a greater proportion of unresponsive cells at day 6 of imaging in WT mice (Extended Data Fig.11D), we looked at the proportions of cell identities regardless of whether neurons were longitudinally tracked or not and found similar proportions of unresponsive cells in WT and CCN1 cKO at day 1 and day 6 of imaging (Extended Data Fig.11J). This suggests some variability in which populations of neurons were longitudinally tracked (Extended Data Fig.11K). In CCN1 cKO mice, we found an increase in binocular cell turnover, with greater monocular to binocular conversion (Extended Data Fig.11H), and no differences in the contralateral or ipsilateral cell turnover (Extended Data Fig.11E and F), indicating that increased binocular cell conversion during the experimental timeline could be inherent to the CCN1 cKO mice independent of manipulation.

Taken together, these data indicate in the absence of astrocyte CCN1 in adulthood, there is both a shift in binocular circuitry, with reduced binocular cells and increased contralateral cells, and a basal increase in binocular cell turnover. Additionally, MD in CCN1 cKO cells leads to a change in the binocular circuitry composition in adults that is absent in the WT. While we find changes in ODI that have been previously reported in the adult in WT mice, namely an increase in nondeprived eye responses (*60*), we do not see these changes in CCN1 cKO mice, suggesting that WT and cKO circuits have different strategies for regulating responsivity after MD. The aberrant binocular connectivity in CCN1 cKO mice could underlie the lack of ODI change in these mice, as perhaps the circuit is already unstable and does not respond to changes in visual input, i.e. monocular deprivation, in the same way that WT circuits do.

### Astrocyte CCN1 regulates binocular circuit properties in adulthood *in vivo*

To examine the role of astrocyte CCN1 in regulating properties of binocular circuits beyond cellular composition, we assessed other key features of these circuits. We looked at orientation tuning for responsive, longitudinally tracked neurons and calculated neurons’ preferred orientations by summating the neural responses to the different orientations across vector space (Methods). We found that CCN1 cKO mice had differences in their preferred orientations for contralateral neurons and for contralateral responses of binocular neurons relative to WT mice (Fig. 5A), with an overall leftward shift towards 0°. However, there were no differences in the proportion of neurons with preferred orientations that were cardinal (Extended Data Fig.12). There were no differences in binocular matching in WT or CCN1 cKO mice before or after MD (Extended Data Fig.10I), in contrast to critical period studies showing that MD does degrade binocular matching (*58*). Moreover, CCN1 cKO mice had lower circular variance of orientation tuning for contralateral cells and contralateral responses from binocular neurons (Fig 5B). This could reflect changes in inhibitory circuit activity in CCN1 cKO mice, as inhibitory circuit activity has been shown to modulate orientation tuning (*61–63*).

**Figure 5.**
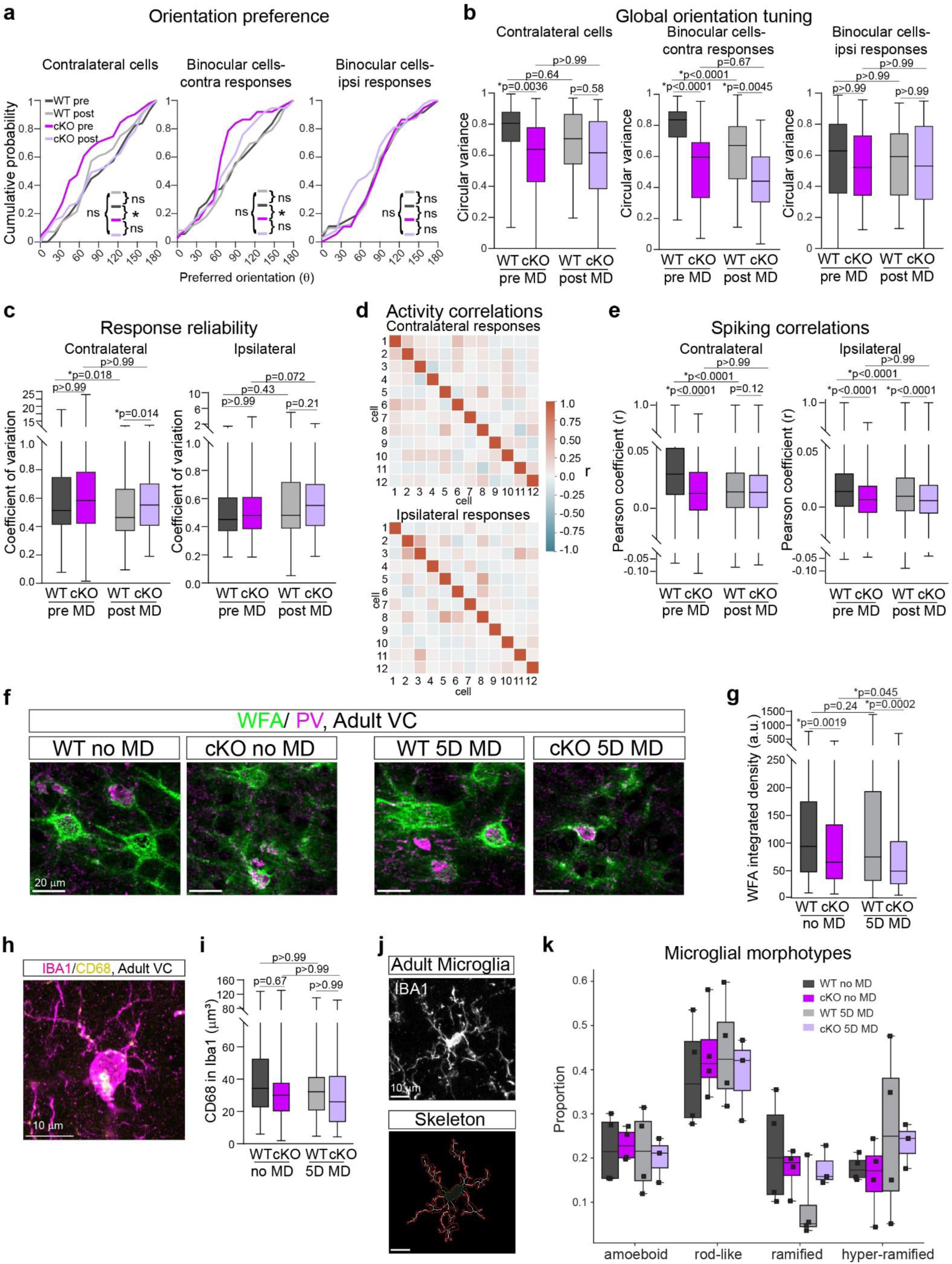
Astrocytic CCN1 regulates binocular circuit properties, maturity of inhibitory circuits, and microglial state in adulthood. **a-b**. Longitudinally tracked contralateral and binocular neurons in WT and CCN1 cKO adult mice. **a**. Orientation preferences. Kolmogorov-Smirnov tests. **b**. Global orientation tuning measured by circular variance. Left to right: contralateral neurons responses, binocular neurons-contralateral responses, binocular neurons-ipsilateral responses. For **a-b**, contralateral neurons: WT = 56, cKO = 51. Binocular neurons: WT = 72, cKO = 37. **c**. Response reliability of all responsive, longitudinally tracked neurons. Left, contralateral eye responses. Right, ipsilateral eye responses. WT = 223 neurons, cKO = 163 neurons. **d**. Representative activity correlation heat map. Scale bar is Pearson coefficient (r). **e**. Spiking correlations of all responsive, longitudinally tracked neurons. Left, contralateral eye responses. Right, ipsilateral eye responses. WT = 9736 pairwise correlations, cKO = 4241 pairwise correlations. **b-e**. Kruskal Wallis tests with post-hoc Dunn’s tests. **f**. Representative images of WFA and PV staining. Scale bar, 20 µm. **g**. WFA integrated density around PV cells. 5 mice per group. WT no MD, n = 235 cells. WT MD, n = 212 cells. cKO no MD, n = 220 cells. cKO MD, n = 198 cells. Kruskal Wallis tests with post-hoc Dunn’s tests. **h**. Representative image of IBA1and CD68 staining in adult VC. Scale bar, 10 µm. **i.** Volume of CD68+ puncta within IBA1 volume. Kruskal Wallis tests with post-hoc Dunn’s tests. **j**. Representative image of IBA1 staining from adult VC, top. Representative cell body and process tracing (skeleton), bottom. Scale bar, 10 µm. WT no MD, 883 microglia from 4 mice. WT MD, 612 microglia from 4 mice. cKO no MD, 667 microglia from 4 mice. cKO MD, 543 microglia from 3 mice. **k**. Proportion of microglial morphotypes-amoeboid, hyper-ramified, rod-like, and ramified. Black symbols, mouse averages. 2-way ANOVA with post-hoc Tukey’s tests. Box charts show median, upper and lower quartiles and whiskers show full range of data.

We also found that astrocytic CCN1 can regulate the response reliability of visual cortex neurons. The coefficient of variation of neuronal responses, a measure of response reliability, to contralaterally presented stimuli decreased in WT mice, but not CCN1 cKO mice, after deprivation (Fig. 5C). This reflects a loss of unreliable contralateral inputs present only in WT mice. Furthermore, looking at pair-wise cellular spiking correlations during stimulus presentation reveals less correlated contralateral responses in CCN1 cKO mice relative to tdT mice (Fig. 5D and E). Contralateral correlations are decreased after MD in both cKO and WT mice. There are no baseline differences in the correlations of ipsilateral responses between WT and cKO mice, though both are decreased after MD (Fig. 5E). Taken together, these results highlight the aberrant binocular circuit properties in CCN1 cKO mice, especially in contralateral eye responses. Thus, astrocyte CCN1 is not only important for maintaining the composition of binocular circuits in adulthood, but their properties as well.

### Astrocyte CCN1 regulates maturity of inhibitory circuits and microglial state in adulthood

Since we found that in critical period age mice, overexpression of CCN1 resulted in increased maturity of inhibitory circuits (Fig. 3), we predicted that loss of CCN1 in adult astrocytes would result in decreased maturity of inhibitory circuits. To address this, we stained for PNNs with WFA as previously described and found that in adults, loss of CCN1 in astrocytes results in decreased PNN density around PV interneurons as well as around all cells (Fig. 5F and G, Extended Data Fig.13A). At baseline, CCN1 cKO mice have decreased PNNs and this is further reduced after 5 days of MD (Fig. 5G), with no change induced by MD in WT mice. Thus, the absence of CCN1 in adult astrocytes results in decreased PNN density, suggesting inhibitory circuits are less mature.

We then assayed microglial interaction with PNNs in adult WT and CCN1 cKO mice as previously described (Extended Data Fig.13B). We found no differences in IBA1-contained CD68 puncta colocalization with WFA across groups (Extended Data Fig.13C) or CD68 volume (Fig. 5H and I). Morphological examination of microglia with clustering analysis (Fig. 5J; Extended Data Fig.14) revealed that at baseline WT mice have a higher proportion of rod-like microglia relative to ramified and hyper-ramified microglia (Extended Data Fig.14B). 5 days of MD in WT mice accentuates these differences and increases rod-like microglia relative to amoeboid. At baseline CCN1 cKO mice have an increased rod-like proportion relative to hyper-ramified and ramified, as in WT mice at baseline. However, 5 days of MD eliminates any significant differences between morphotypes (Fig. 5K; Extended Data Fig.14B); this was supported by an increase in the total skeleton length and a decrease in form factor (Extended Data Fig.14C-F). Rod-like microglia have typically been described in aging, neuronal injury, and age-associated diseases (*64*), but they could also reflect a transition state to activated morphotypes (*49*). While the effects of CCN1 and monocular deprivation on microglial morphotype are less dramatic in adult mice than in critical period aged mice, they nevertheless demonstrate the role of CCN1 in regulating microglial state in an age-specific and context-specific manner.

## Discussion

Our studies show that astrocytes induce synaptic stability and regulate circuit connectivity in the adult visual cortex. Stemming from an RNA sequencing screen, we establish CCN1 as an astrocyte-secreted factor whose expression increases in adulthood and is downregulated when plasticity is induced (Fig. 1). We show that prematurely upregulating CCN1 in critical period aged mice restricts binocular zone remodeling and drives the maturation of inhibitory circuits, shown by increased excitatory synaptic drive onto fast-spiking interneurons and increased PNN density (Fig. 2 and 3). Conversely, the absence of CCN1 in adult astrocytes enables enhanced binocular zone remodeling and decreases the maturation state of inhibitory circuits, as evidenced by decreased PNN density in CCN1 cKO mice (Fig. 2 and 5). The bi-directional impact of astrocytic CCN1 on PNN levels has implications for tuning inhibitory circuit function and overall network activity (*65, 66*).

Our studies also demonstrate that visual circuits, even after the establishment of binocularity during early visual experience and the critical period (*56, 57*), require ongoing maintenance cues, and that astrocytes provide these cues via the secretion of CCN1. This is evident in adult CCN1 cKO mice, where its absence results in decreased numbers of binocular responsive neurons and increased numbers of contralateral responsive neurons. As our data show that adult WT mice undergo neuronal response magnitude changes but cKO mice undergo binocular circuitry composition changes after MD, this suggests that in the absence of astrocytic CCN1 different plasticity rules are engaged (Fig. 4). Moreover, binocular circuits in CCN1 cKO mice show differences in global orientation tuning, response reliability, and activity correlations, demonstrating that CCN1 from astrocytes can regulate the functional properties of binocular circuits (Fig. 5). The changes in inhibitory circuit maturation we found could be a mechanism underlying these changing plasticity rules and properties (*67*).

A mechanism by which CCN1 regulates visual circuit synapse stability may be through signaling to microglia. Microglia have been established as important regulators of the extracellular matrix (*46, 48, 68–70*) and synapse number (*46, 68, 70–75*) through engulfment and phagocytosis in plasticity and disease models. Our results show that CCN1 from astrocytes modulates microglial morphology (Fig. 3 and 5) in the visual cortex. We found that CCN1 overexpression in critical period age mice results in an increase in the proportion of ramified microglia and a decrease in amoeboid microglia after monocular deprivation relative to control mice. Amoeboid morphotypes are typically more associated with phagocytosis and a pro-inflammatory profile (*52*). These data, in conjunction with the reduction of phagocytotic lysosomes in CCN1 o/e mice, suggest that CCN1 from astrocytes during the critical period may be resulting in impairments in phagocytosis and microglial activation. The changes to microglia in adult CCN1 cKO mice are less pronounced, though there are changes in the proportion of rod-like microglia (Fig. 5), a morphotype associated with aging and neuronal injury. The critical period and adult data suggest that bidirectionally regulating the expression of CCN1 in astrocytes can affect microglial state in a context-specific manner. Since CCN1 has been shown to be upregulated in stroke (*76*), and is a putative immediate early gene in astrocytes (*77*), further experiments should interrogate whether astrocytic CCN1 can impact microglial engulfment of synapses or PNNs in disease or acute injury models.

In conclusion, our studies establish the crucial role of astrocytes in actively stabilizing the connectivity of neuronal circuits. They highlight the impact that astrocytes have on different cell types in the brain, including neuronal and non-neuronal cells. In turn, these findings deepen our understanding of how stability of sensory circuits is actively maintained in the adult brain, through a complex interaction of multiple cell types. Understanding what makes the neural environment permissive to plasticity in young animals and restrictive in older animals will open new avenues for therapies for recovery after brain injury such as stroke.

## Methods

### Mice

All animal experiments were approved by the Salk Institute Institutional Animal Care and Use Committee (IACUC). Mice were typically housed with a 12-hour light/dark cycle in the Salk Institute animal facilities. Dark reared mice were housed in 24-hour dark cycle since birth. Mice were provided access to food and water *ad libitum*. For transcriptomic experiments, astrocyte-Ribotag mice were generated by crossing GFAP-cre hemizygous females (B6.Cg-Tg (GFAP-cre)73.12Mvs/J, Jax #012886) to homozygous flox-Rpl22-HA males (B6N.129-Rpl22tm1.1Psam/J, Jax #011029). Male mice hemizygous for Cre and heterozygous for flox-Rpl22-HA (Rpl22-HA+; GFAPcre+) were used for all experiments. Wild-type C57Bl6/J mice were used (Jax 000664) for experiments shown in Figures 1-3. For single molecular fluorescence *in situ* hybridization experiments (Fig. 1H) validating the transcriptomics, male mice were used. For adult knock-out experiments, CCN1^fl/fl^ mice were a gift from Dr. Lester Lau (*36*) and were maintained on a C57Bl6/J background. These mice were crossed to mice expressing tamoxifen inducible Cre recombinase under an astrocytic promoter for temporal elimination (Aldh1l1cre-Ert2, Jax 029655(*78*)). Experimental mice were homozygous for the CCN1 floxed allele and either Cre- or Cre+ (WT or CCN1 cKO). Mice were injected intraperitoneally (i.p.) with 75 mg/kg of tamoxifen (MP Biomedicals #156738) for 5 consecutive days at 1 month of age. Mice of both sexes were used and sexes were noted.

### Surgical procedures

#### Juvenile viral injections

For AAV injections at postnatal day 14-15 (P14-15), C57Bl6/J mice were used. Briefly, mice were administered pre-operative carprofen (5 mg/kg) subcutaneously and anesthetized using isoflurane. Stereotaxic coordinates for the binocular zone (BZ) were 2.25 mm lateral and 0.5 mm anterior from lambda. Virus was injected at 3 sites at a depth of 500-600 µm from just below the skull surface. The pipette was kept in the brain for 3 minutes after each injection to allow the virus to diffuse. Both tdT and CCN1 viruses were injected for a total titer of ∼2 x 10^8^ vg/mL. After injection, mice were sutured and placed back with the dam.

#### Monocular enucleation and monocular deprivation

For monocular enucleation at P28 or 4 months, mice were anesthetized using isoflurane. The eye was removed using curved forceps and pressure was applied to stop any bleeding. GelFoam was inserted into the eye socket and 2 box sutures (Henry Schein 5616446) were used to close the eyelid. Mice were monitored daily and administered ibuprofen water (0.15 mg/ mL) to minimize any swelling.

For monocular deprivation at P28 or 4 months, mice were anesthetized using isoflurane. Eyelashes were trimmed down to the eyelid margins and 4 box sutures using nylon sutures (Ethilon 1647G) were used to close the eyelid. Mice were monitored daily and administered ibuprofen water (0.15 mg/ mL) to minimize any swelling. Mice were removed from the experiment if the eyelids opened. For suture removal, mice were again placed under isoflurane and sutures were removed. Mice were removed from the experiment if the eye looked damaged or cloudy.

#### Cranial window implantation and viral injections

For *in vivo* imaging of ocular dominance plasticity and neuronal response properties, cranial windows were implanted on ∼3 month old CCN1 WT or cKO mice. Mice were injected with Carprofen (5 mg/kg, s.c.), Buprenorphine SR (1 mg/kg, s.c.), Baytril (10 mg/kg, i.m.), and Dexamethasone (2 mg/kg, i.p.) prior to surgery for anesthesia, infection prevention, and inflammation prevention. Mice were anesthetized with isoflurane inhalant (3%) and maintained at 1.5-2.5% during the surgery. Mice were mounted on a stereotaxic surgical stage via ear bars and a bite bar. Their body temperature was maintained at 37°C using a heating pad. The scalp was shaved and the skin was removed and the skull surface was allowed to dry. The skull and scalp margins were covered with a thin layer of Vetbond (Fisher Scientific NC0304169). Avoiding the area above visual cortex, a layer of dental cement (Tetric evoflow A1, Henry Schein 9458634) was applied and a metal head plate was glued to skull. A 3 mm circular piece of skull over the binocular zone (coordinates from lambda: 3 mm lateral, 1.0 mm anterior) was removed with a high speed microdrill with a 0.5 mm burr. Care was taken not to damage the dura. Before window implantation, viral injections of AAV2/1-CAMK1a-gCaMP6f (Addgene #100834-AAV1) were made into the binocular zone. 150 nL was injected into each of 5 sites at a depth of 300 µm from the pia for a total titer of ∼1.2 x 10^10^ vg. The pipette was kept in the brain after each injection for 3 minutes to ensure diffusion of the virus before being retracted. A 4 mm diameter coverslip was placed on the dura and sealed to the edges of the skull using Vetbond. Dental cement was then used to further seal the coverslip. Mice were injected subcutaneously with 1 mg/ kg physiological saline and placed on a heating pad to recover. Carprofen (5 mg/kg, s.c) was administered daily for 3 days post-surgery. Mice were maintained with Baytril (8.5 mg/kg/ day) in their water to prevent infection for 3 days.

### RNA sequencing

Data for experiments from P28 (critical period) and P120 (adult) mice were obtained from Farhy-Tselnicker et al. 2021 (*19*) (GEO GSE161398) and Boisvert et al. 2018 (*28*) (GEO GSE99791).

#### Conditions for analysis

##### Developmental time course

For experiments comparing mice P28 (critical period) to P120 (adult), see Farhy-Tselnicker et al. 2021 (*19*) and Boisvert et al. 2018 (28) for collection methods.

##### Dark rearing

Dark rearing (DR) was performed by housing mice in ventilated telemetry cabinets, in complete darkness, and all husbandry and cage changes were done under red light. Mice for dark rearing were born in the dark and remained there until P45, anesthetized under red light, and perfused with a hood over their head to prevent light from reaching the eyes. For age-matched comparison mice were raised under 12hour light/12hour dark cycle until P45. The visual cortices from 2 mice (Rpl22-HA+; GFAPcre+) were pooled for RNA isolation and RNA sequencing library preparation. P45 DR = 4 biological replicates (8 mice, 2 x 4). P45 = 3 biological replicates (6 mice, 2 x 3).

##### Monocular Deprivation

Monocular deprivation (MD) was performed at P26, for 2 days until P28, during the peak of the critical period. The visual cortex contralateral to the deprived eye (major loss of visual input) and ipsilateral to the deprived eye (minor loss of visual input) were collected separately for analysis and comparison. The visual cortices from 2 mice (Rpl22-HA+; GFAPcre+) were pooled for RNA isolation and RNA sequencing library preparation. MD contra = 3 biological replicates, (contralateral visual cortex from 6 mice, 2 x 3); MD ipsi = 3 biological replicates, (ipsilateral visual cortex from 6 mice, 2 x 3).

### Ribotag Pulldown and RNA sequencing

Male mice heterozygous for flox-Rpl22-HA (Jax 011029) and hemizygous for GFAP-cre (Jax 012886) (astrocyte-ribotag) were used to isolated astrocyte mRNA based on a modified ribotag protocol (*79*).

#### Dissection

All mice were collected between 9:30am and 12:30pm on the day of experiment. Dissection was performed as described in Farhy-Tselnicker et al. 2021 (*19*) and Boisvert et al. 2018 (28).

#### Ribotag pulldown

A modified Ribotag protocol was performed as described in Farhy-Tselnicker et al. 2021 (*19*) and Boisvert et al. 2018 (28).

#### RNA sequencing library generation and sequencing

Library preparation was performed as described in Farhy-Tselnicker et al. 2021 (*19*) and Boisvert et al. 2018 (28).

#### RNA sequencing mapping, analysis, and statistics

Sequencing data mapping, analysis, and statistics were done as described in Farhy-Tselnicker et al. 2021 (*19*) and Boisvert et al. 2018 (28).

### Selection of differentially expressed genes (DEGs)

1. FPKM > 1 in each sample of at least one group per comparison
2. Ribotag pulldown FPKM (astrocyte)/input FPKM (all cells) > 0.75 in each at least one group per comparison
3. Adjusted p-value < 0.05
4. Log2 Fold Change (LFC) > |0|

Venn diagram of overlapping DEGs in different experimental comparisons were generated using R. Heatmaps showing the LFC of DEGs were generated in R using the Pheatmap package. Predicted functional interaction network of CCN1 was generated using the STRING database (*80*) (Fig. 1).

#### Pathway analysis

Gene-set enrichment analysis (GSEA) was performed to determine which predefined sets of genes were significantly enriched across the plasticity paradigms (*81*). Enrichment of gene sets and pathways from the Gene Ontology (GO), Reactome, KEGG and WikiPathway databases was carried out. Only genes that had an FPKM > 1 in each sample were included. Genes were ranked based on descending log2 Fold Change (LFC) in each comparison. A cutoff of adjusted p value < 0.05 was used to determine significantly enriched pathways and terms. Simplifying GSEA results for visualization was conducted by selecting the top three down- and upregulated (negative and positive normalized enrichment score (NES), respectively) Reactome pathways in each comparison (Extended Data Fig.1B). Overrepresentation analysis (ORA) was also performed to determine GO terms that were enriched in the significant DEGs across the plasticity paradigms (Extended Data Fig.2). Lists of upregulated and downregulated DEGs based on the same criteria as for the heatmaps and Venn diagrams were included and the background list of genes were those that had FPKM > 1 in each sample of at least one group per comparison. The following criteria were used to select significantly enriched GO terms: adjusted p value < 0.05 (Benjamini and Hochberg correction) and q-value < 0.01. The clusterProfiler package (version 4.10.0) was used to perform GSEA and ORA (*82*).

### Cloning

For the overexpression of HA-tagged CCN1, cDNA for the coding sequence of mouse CCN1 (Origene # MR221828) was used. This cDNA was myc-Flag tagged, so we amplified the cDNA with PCR primers (forward: GCGATCGCCATGAGCTCC, reverse: ttaaccggttgcataatccggaacatcatacggataGAGCGGCCGCGTACG) to insert an HA-tag at the end of the c-terminus in lieu of myc-Flag. The fragment was then amplified again with primers optimized for InFusion cloning (Takara Bio #638909) (forward: cgactcactataggctagcgccaccATGAGC, reverse: tgtctgctcgaagcggccgcttaaccggttgcataatccggaacatcatacg). The linearized product was run out on a 0.8% agarose TAE gel and extracted. We used a pZac2.1 AAV2 backbone for overexpression. pZac2.1-gfaABC1D-tdTomato (GFAP-tdTomato) from Addgene (#44332) was digested with NheI and NotI to remove the tdTomato. The linearized product was run out on a 0.8% agarose TAE gel and extracted. An InFusion reaction was performed with the linearized PCR-amplified CCN1-HA and the linearized vector. Clones were selected using carbenicillin and sequenced to confirm presence of the inserted CCN1-HA. The cloned CCN1 plasmid was validated *in vitro* using astrocyte and HEK 293T/Cre cell cultures. The GFAP-CCN1-HA plasmid was sequenced and sent to the Salk Institute Gene Transfer, Targeting, and Therapeutics Core and was packaged into an AAV2/5 virus. Obtained titer ranged from 2-4 x 10^12^ vg/mL. For the control virus, we obtained the AAV2/5 virus of the GFAP -tdTomato vector from Addgene (#44332-AAV2/5). Titers ranged from 1-4 x 10^13^ vg/mL.

### Cell culture

For validation of the plasmids, *in vitro* over-expression was performed in astrocytes purified from Sprague-Dawley P1-2 rats and in human embryonic kidney (HEK) 293T/Cre cells. The astrocytes were purified using the McCarthy-de Vellis method (MD) (*33*), and grown in culture media containing DMEM (Life Technologies #11960044), 10% Fetal bovine serum (Life Technologies #10437028), 1% Penicillin-Streptomycin (Life Technologies #15140122), 1% Glutamax (Life Technologies #35050061), 1% sodium pyruvate (Life Technologies #11360070), 5 ug/mL NAC (Sigma # A8199), 5 ug/mL insulin (Sigma #11882), and 10 µM hydrocortisone (filtered with 0.22 µm filter). Cell culture dishes were coated with poly-D-lysine (Sigma #P6407) before splitting MD astrocytes onto them. MD astrocytes were passaged 2-3 times before being transfected.

HEK cells were grown in HEK cell growth media containing DMEM, 10% fetal bovine serum, 1% Penicillin-Streptomycin, 1% Glutamax, and 1% sodium pyruvate. HEK cells were passaged 4-5 before being transfected.

#### Plasmid validation

To validate plasmids via immunocytochemistry (ICC) (Extended Data Fig.4A and B), cultured astrocytes and HEK cells were plated in 24 well plates containing glass coverslips. For HEK cells, coverslips were coated with 1:50 CELLstart (ThermoFisher #A1014201) diluted in water. For astrocytes, coverslips were coated with poly-D-lysine as described above. Both HEK cells and astrocytes were transfected the day after plating with 500 ng of plasmid DNA using lipofectamine 2000 (Invitrogen #11668019) and OptiMEM (Life Technologies #31985-070). After transfection, astrocytes were maintained in astrocyte growth media, while HEK cells were maintained in HEK cell growth media. After 5 days of expression, cells were fixed with 4% PFA. Cells were then permeabilized with 1% BSA and 0.2% Triton X-100. Coverslips were incubated overnight at 4C in primary antibodies diluted in 1% BSA. For HEK cells, rabbit anti-HA antibodies (CST #3724), and sheep anti-CCN1 (R&D System AF4055) were used at 1:500. Secondaries were used at 1:1000 for 2 hours at room temperature. For astrocytes, mouse anti-GFAP (Millipore #360) and rabbit anti-HA were used. SlowFade Gold with DAPI mounting media (LifeTech #S36939) was used. Coverslips were imaged using an Axio Imager.Z2 fluorescent microscope (Zeiss) with an AxioCam HR3 camera (Zeiss) at 20x magnification.

### Mouse tissue collection

Tissue for single-molecule fluorescent *in situ* hybridization (smFISH) in Fig. 1 and Extended Data Fig.3 was collected at P7, P14, P28, P45, and P120. Tissue for CCN1 cKO validation was collected at 2 months of age. Tissue for smFISH against Arc was collected at P33 or 4 months of age. Tissue for immunohistochemistry was collected at approximately 1 month of age and 4 months of age.

#### smFISH

All mice for smFISH were collected between 1pm and 5pm. Mice were anesthetized by i.p. injection of 100 mg/kg ketamine (Victor Medical Company)/20 mg/kg xylazine (Anased) mix and transcardially perfused with PBS. Brains were removed and embedded in OCT media (Sakura 4583), frozen in dry ice-ethanol slurry solution, and stored at –80°C until use. Sagittal sections were obtained using a cryostat (Hacker Industries #OTF5000) at a slice thickness of 16–20 µm. Sections were mounted on Superfrost Plus slides (Fisher #1255015). smFISH was performed on the same day as sectioning. 3–6 mice were used for each experimental group. For each mouse, 2-3 sections were imaged and analyzed.

#### *Arc* induction

All mice for *Arc* induction were collected from the vivarium between 5:00-5:30am at the end of the mice’s dark cycle. Mice were brought back to the lab space and exposed to bright light for 30 minutes. Mice were then anesthetized by i.p. injection of 100 mg/kg ketamine (Victor Medical Company)/20 mg/kg xylazine (Anased) mix. Mice were immediately decapitated, brains were extracted, embedded in OCT and flash frozen in dry ice-ethanol slurry mix. Brains were stored at -80C until use. Coronal sections were obtained using a cryostat at a slice thickness of 16–20 µm. Sections were mounted on Superfrost Plus slides (Fisher #1255015). smFISH was performed on the same day as sectioning. 4–6 mice were used for each experimental group. For each mouse, 4-6 sections were imaged and analyzed.

#### Immunohistochemistry

Mice were anesthetized by i.p. injection of 100 mg/kg ketamine /20 mg/kg xylazine mix and transcardially perfused with phosphate buffered saline (PBS), then 4% PFA at room temperature. Brains were removed and incubated in 4% PFA overnight at 4°C, then washed 3 x 10 minutes with PBS, and cryoprotected in 30% sucrose for 2–3 days. Brains were then embedded in tissue freezing media (TFM; General Data Healthcare #TFM-5), frozen in dry ice-ethanol slurry solution, and stored at –80°C until use. Brains were sectioned using a cryostat (Hacker Industries #OTF5000) in sagittal or coronal orientations depending on experimental needs at a slice thickness of 16–20 µm. Sections were mounted on Superfrost Plus slides (Fisher #1255015). 3–6 mice were used for each experimental group. For each mouse, 2-3 sections were imaged and analyzed.

### Histology

#### Immunohistochemistry on mouse brain tissue

The slides containing the sections were blocked for 1 hr at room temperature in blocking buffer consisting of 1% BSA and 0.2% Triton X-100 diluted in PBS. Primary antibodies were diluted in this blocking buffer and incubated overnight at 4°C. The next day, slides were washed 3 x 10 minutes with PBS and secondary antibodies conjugated to Alexa Fluor (Molecular Probes) were diluted in blocking buffer and applied for 2 hrs at room temperature. Slides were mounted with the SlowFade Gold with DAPI mounting media, covered with 1.5 glass coverslip (Fisher #12544E), and sealed with clear nail polish.

For validation of the viral vectors in Fig. 2 and Extended Data Fig.4C, the following antibodies were used: goat anti-SOX9 (R&D Systems af3075, 1:250), rabbit anti-HA (CST #3724, 1:500), mouse anti-NEUN (Millipore #MAB377 1:100), mouse anti-GFAP (Millipore #360, 1:500). Secondary antibodies were donkey anti-goat Alexa Fluor 488 (Jackson ImmunoResearch 705-545-147), donkey anti-rabbit Alexa Fluor 568 (ThermoFisher A-10042), donkey anti-mouse Alexa Fluor 647 (Jackson ImmunoResearch 715-605-150), goat anti-mouse Alexa Fluor 647 (ThermoFisher A-21235). All secondary antibodies were applied at 1:500 dilution.

For perineuronal net (PNN) deposition experiments, the following antibodies were used: rabbit anti-HA (CST #3724, 1:500), mouse anti-parvalbumin (Millipore Sigma p3088, 1:500). Biotinylated wisteria floribunda agglutinin (WFA, Vector Laboratories B-1355-2, 1:500) was used at the same time as the primary antibodies to stain for the PNNs. Secondaries were used at 1:500 and included goat anti-rabbit Alexa Fluor 647 (ThermoFisher A-21245), goat anti-mouse Alexa Fluor 488 (ThermoFisher A-11001) and streptavidin conjugated Alexa Fluor 568 (adult mice, ThermoFisher S-11226) or 647 (critical period mice, ThermoFisher S-21374) to stain for PNNs.

For microglia morphology and phagocytosis experiments, the following antibodies were used: rabbit anti-IBA1 (Fuji Film Wako #162001, 1:500), rat anti-CD68 (BioRad #MCA1957GA, 1:100), and biotinylated WFA (Vector Laboratories B-1355-2, 1:500). To confirm for viral expression in critical period aged mice, half the sections on each slide were stained with rabbit-anti HA (CST #3724, 1:500) instead of staining for microglia morphology and engulfment. Secondaries were used at 1:500 and were goat anti-rabbit Alexa Fluor 488 (ThermoFisher A-11008), goat anti-rat Alexa Fluor 594 (ThermoFisher A-11007), and streptavidin conjugated Alexa Fluor 647 (ThermoFisher S-21374).

#### Single-molecule fluorescent in situ hybridization (smFISH)

All smFISH experiments were done as described in Farhy-Tselnicker et al. 2021 (19) except on fresh-frozen tissue as described in the “Mouse tissue collection” section. For tissue from P7 mice, slides were incubated with protease plus for 15 min; for P14–P120, protease 4 10-20 min. Detailed step-by-step modified protocol performed here is available upon request. Probes used were: Ccn1 (ACDbio #429001), Tubb3 (423391-C3), Slc1a3 (430781-C2).

### Imaging and analysis

#### Epi-fluorescence microscopy

Imaging was performed using an Axio Imager.Z2 fluorescent microscope (Zeiss) with the apotome module (Apotome2) and AxioCam HR3 camera (Zeiss) at 10x or 20x magnification, depending on the experiment. Tile images that contained the entire primary visual cortex (from pial surface to white matter tract) were acquired. Number of tiles were maintained consistent within each experiment. Images were 14 bit, All tiles had 10% overlap. For critical period experiments looking at WFA expression, all sections were confirmed to have viral vector expression.

For the Arc smFISH (Fig. 2), images were acquired at a single plane at 10x, and the entire section was tiled.

For smFISH in Fig. 2 and Extended Data Fig.4E for the CCN1 cKO validation, images were acquired at 20x, with 3 tiles of the entire visual cortex taken, and a z-stack width of 10-12 um.

For the immunohistochemistry for viral vector validation in Fig. 2, images were acquired at 20x with 6 tiles taken of the visual cortex at 10% overlap. For WFA/ PV staining, images were acquired at 20x with 3 tiles taken of the visual cortex at 10% overlap.

In all cases, when comparing WT and cKO or different viruses per given experiment, slides were imaged on the same day using the same acquisition settings.

#### Confocal and super-resolution microscopy

smFISH images used for layer-specific developmental profile of *Ccn1* expression were acquired using a Zeiss LSM 700 confocal scanning microscope. Images were acquired at 8-bit depth, 1024 x 1024 resolution using a 20x objective with a pixel dwell time of 0.79 µs. The scaling per pixel was 0.31 µm x 0.31 µm and 2 frames were averaged per plane. Z-stacks at 1 µm steps were taken and the visual cortex was tiled at 10% overlap. Number of tiles remained consistent between experiments.

For microglia engulfment analysis, a Zeiss LSM 880 with Airyscan module was used. The images were acquired in super-resolution mode with a Fast Airyscan module using an oil-immersion 63x objective with a numerical aperture of 1.46 at a pixel dwell time of 0.60 µs. The scaling per pixel was 0.041 µm x 0.041 µm. Z-stacks with 0.110 µm steps were acquired using a piezo, typically 150-200 planes per image. For the critical period mice expressing tdTomato, the 563 laser was used to photobleach the tdTomato signal prior to imaging the engulfment. To confirm that critical period mice injected with CCN1-HA, we confirmed viral vector expression with half the sections on each slide that were stained with rabbit-anti HA. The WFA channel (647) was imaged with an excitation beamsplitter of 488/561/633 and an emission filter set of 570-620 bandpass and 645 longpass. The CD68 channel (594) was imaged with an excitation beamsplitter of 488/594 and an emission filter set of 420-480 bandpass and 495-620 bandpass. The IBA1 channel (488) was imaged with an excitation beamsplitter of 488/561/633 and an emission filter set of 420-480 bandpass and 495-550 bandpass. Laser power for acquisition was kept the same across experiments. Images were Airyscan processed in automatic mode.

For microglia morphology analysis, the same microscope was used but in normal confocal mode. Images were acquired at 16-bit depth using a 20x objective with a pixel dwell time of 2.05 µs. The scaling per pixel was 0.21 µm x 0.21 µm. Z-stacks at 2 µm steps were taken and the visual cortex was tiled at 15% overlap. Number of tiles remained consistent between experiments.

#### Image analysis

All image analysis of the smFISH in Fig. 1 and the CCN1 cKO validation in Figure 2 was performed in ImageJ using a custom macro developed in the lab. The *Slc1a3* signal was thresholded and used to segment astrocyte regions of interest (ROI). The *Ccn1* probe signal was also thresholded and the total area within each astrocyte ROI was quantified. Thresholds were kept the same within experiments (e.g. ages).

For quantification of the IHC validating the viral vectors, a Cell Counter plug-in in ImageJ was used to count the number of viral transduced cells. For quantifying astrocyte GFAP level in Extended Data Fig.4C and D, GFAP signal was thresholded equally in each pair of mice (tdT vs CCN1) and total area was quantified and compared.

To quantify the level of the perineuronal net (PNN) around parvalbumin (PV) cells, a custom region of interest (ROI) detection pipeline was developed in CellProfiler. The PV channel intensity was rescaled in order to use the full intensity range to increase the brightness of the image for ROI selection. Both channels had a gaussian filter with a sigma of 1 applied. In order to reduce image heterogeneity and optimize ROI detection, the lower quartile of pixel intensity values were subtracted from the WFA and PV channels. ROIs were manually curated and added/removed. Finalized WFA and PV ROIs were then overlayed with the original, unprocessed images and the integrated density of WFA or PV per ROI was obtained. To separately quantify only WFA that surrounded PV cells, ROIs were overlayed and excluded if overlap was absent. The overlapping ROIs were overlayed with the original, unprocessed images and the integrated density per ROI was obtained. Values were exported to a csv file.

For analysis of microglia engulfment, images were analyzed using pyclesperanto in a Napari viewer (https://github.com/clEsperanto/napari_pyclesperanto_assistant). Briefly, the WFA, IBA1, and CD68 channels were independently segmented. A gaussian filter followed by a gamma correction at 1.5 were applied to the WFA signal prior to thresholding using the Otsu method. The IBA1 and CD68 signals were thresholded using the Otsu method. However, for critical period mice, for the CD68 signal the manual thresholding value was set to 900 instead of using the Otsu method due to differences in the CD68 signal in critical period aged mice. The overlapping regions (WFA+CD68+IBA1 and CD68+IBA1) were generated using the segmentations and saved as tiffs. To extract volumes of the segmentations, the tiffs were analyzed in ImageJ using the Analyze 3D objects plugin and using the same thresholding for all images (Extended Data Fig.7 and 13).

#### Microglia morphological classification

For microglia morphology analysis, images were processed in a custom ImageJ macro and analyzed using a custom CellProfiler pipeline. In ImageJ, the contrast for each image was normalized so that at least 70% of the pixels are saturated, and the image was despeckled to remove noise. The pia and white matter were excluded from analysis to standardize the microglia morphology in each image. Then, in CellProfiler, the EnhanceOrSuppressFeatures module was used to suppress feature sizes of 10 to segment soma from processes. The soma was identified as the primary object via the adaptive two-class Otsu thresholding method (threshold smoothing scale: 1, threshold correction factor: 1, size of adaptive window: 100). Soma area was measured using the MeasureObjectSizeShape module. The non-suppressed image underwent a Gaussian filter (sigma 1) and thresholded using the global robust background method (averaging method: mean, variance method: standard deviation, # of deviations: 0.8, threshold smoothing scale: 1, threshold correction factor: 0.9). From the thresholded image, secondary objects (processes) were identified via propagation from the overlaid primary objects (soma) using the adaptive two-class Otsu thresholding method (threshold smoothing scale: 0.3, threshold correction factor: 1, size of adaptive window: 10, regularization factor: 1). Microglia that had processes touching the borders of the image were excluded from analysis to remove artificial decreases in measurements. The MorphologicalSkeleton module was used to generate skeletons within the processes, and the MeasureObjectSkeleton module quantified number of trunks, non-truck branches, and total skeleton length of each microglia. Average branch length was calculated by dividing the total skeleton length by the sum of the number of trunk and non-trunk branches. All measurements were converted to microns.

For microglial morphology clustering analysis, Python sci-kit learn was used to scale and perform dimensionality reduction on all the extracted morphology features using principal component analysis (PCA). The number of components was chosen as the minimum number that explained 80% of the variance. Then, k-means clustering was performed and the elbow method was used to minimize the within cluster sum of squares. The number of chosen clusters was 4 for both critical period and adult mice, which is in line with the literature (*49–52*). To plot the cluster heatmaps, we used the scaled mean of each feature; we identified each cluster as amoeboid, rod-like, ramified (resting), or hyper-ramified based on descriptions from the literature. The mean proportion per cluster for each mouse was calculated and a linear mixed effects model with a beta distribution was built in R using the glmmTMB and DHARMa libraries (*83*). Posthoc tests looking at pairwise comparisons of clusters across viral groups/ genotypes and manipulations were performed with p-value adjustments for multiple comparisons.

### Electrophysiology

#### Acute slice preparation

Coronal slices of the binocular zone of the visual cortex were prepared from P32-33 WT mice, that had been injected at P14 with GFAP-TdTomato or CCN1-HA, following protocols in Kuhlman et al., 2010 and Xue, Atallah and Scanziani, 2014. Animals were deeply anesthetized by injection with Avertin and immediately decapitated. The brain was removed, hemi-sected, and cut into 300 µm coronal sections using a Leica VT1000s vibratome. The brain dissection was performed in cold, sucrose-based dissection solution consisting of (in mM): 2.5 KCl, 7.0 MgCl2, 1.25 Na2HPO4, 11 glucose, 234 sucrose, 0.50 CaCl2, and 24 NaHCO3 and equilibrated with carbogen (95% O2/ 5% CO2). Slices were then placed in a recovery chamber containing artificial cerebrospinal fluid (aCSF) consisting of (in mM): 126 NaCl, 26 NaHCO3, 1.25 Na2HPO4, 2.5 KCl, 2 CaCl2, 1 MgCl2, 25 glucose, and saturated with carbogen. Slices recovered for 30 min at 34°C and were then maintained at room temperature until recordings were performed for 4-6 hours after slicing.

#### Electrophysiology

Slices were placed in a recording chamber and perfused with a recirculating bath of carbogen-saturated aCSF maintained at 31°C. Whole-cell patch clamp recordings were obtained from neurons in layer II/III of V1 that were visualized using IR-DIC on a Scientifica microscope. Open pipette resistances were 2-5 MΩ (borosilicate glass pipette; Harvard Apparatus #30-0057). Recordings were performed using a Multiclamp 700B amplifier (Molecular Devices). All recordings were sampled at 10 kHz. For measuring the cell membrane properties, data were filtered at 10kHz. Recordings were discarded if the series resistances were > 25 MΩ or changed > 25% during the entire recording. For recording spontaneous excitatory postsynaptic currents (sEPSCs) in voltage clamp, aCSF containing 40 µM bicuculline methochloride (Toris #0131) was washed in. sEPSCs were filtered at 2 kHz and acquired at a gain of 10 to allow for detection of small events. Fast-spiking cells were confirmed as fast-spiking with little-to-no spike-frequency adaptation by injecting a 100 ms current step and noting the frequency and AP width in current-clamp mode. Additionally, fast-spiking neurons had a low input resistance (<150 MΩ) and multipolar morphology (*38*).

In all experiments, a potassium (K)-gluconate internal consisting of (in mM) was used: 115.0 K-gluconate, 20.0 KCl, 10.0 phosphocreatine disodium salt, 10.0 HEPES acid, 0.2 EGTA, 4.0 Mg-ATP, and 0.3 Na-GTP. Osmolarity and pH of the internal solutions were adjusted to 290-310 mOsm and pH 7.3-7.4 with double-distilled water and with KOH.

#### Data analysis

Off-line data analyses were performed using MiniAnalysis (Synaptosoft) and MATLAB. sEPSC recordings were filtered post-hoc with a 1 kHz low-pass Bessel 8-pole filter in Clampfit (Molecular Devices).

### *In vivo* two photon imaging

#### Two-photon calcium imaging

After recovery from viral injection and cranial window implantation, mice were habituated to handling and head fixation on a linear treadmill (LabMaker) over the course of one week. We used a custom-built two-photon laser scanning microscope from Neurolabware equipped with a pulsed femtosecond Ti:Sapphire laser (Chameleon Vision II, Coherent) controlled by Scanbox acquisition software. The laser was tuned to 920 nm for imaging gCaMP6f and focused through a 16 x water-immersion objective (Nikon, 0.8 numerical aperture). Images were acquired at a frequency of 15.5 Hz at a depth of 150-300 µm below the pial surface for layer 2/3. Images were 512 x 796 pixels (500 x 600 µm). A rotary encoder was used to track the running speed of the mice on the linear treadmill. A camera fitted with a 740 nm long-pass filter was used to track pupil diameter during imaging (Mako U-015B). Both the encoder and the pupil camera were triggered at the scanning frame rate. To measure the responses of neurons to presentation of visual stimuli to either eye independently, an opaque eye path was placed in front of the other, non-imaged eye.

#### Binocular zone confirmation

Imaged areas that were stereotaxically identified during cranial window implantation (see above) as the binocular zone (BZ) were confirmed as such by performing retinotopic mapping. A BENQ LCD 27” monitor (60 Hz refresh rate) was used for visual stimulus presentation, was placed 20 cm from the eyes, and covered approximately 113° in azimuth and 60° in elevation. Stimuli were designed in PsychoPy 2.22 and custom code was written to enable communication between the stimulus computer and the imaging computer. Stimulus presentation onset was tracked using a photodiode (Thorlabs, FDS1010) that was fed to an Arduino UNO Rev3 and generated a transistor-transistor logic (TTL) pulse that was sampled by the imaging computer and time-stamped with the imaged frame. To confirm that the imaged area was indeed binocular, small checkerboard stimuli were presented at 15 neighboring positions at a rate of 3 Hz. Stimuli were presented to either eye independently and regions were considered binocular if the peak responses in both eyes were evoked by stimuli presented in the central, upper visual field (-20° to +20° azimuth relative to the midline). Regions with no or weak ipsilateral responses were also not considered binocular.

#### Visual stimulus presentation for ocular dominance mapping

Drifting sinusoidal gratings were presented to each eye independently. The stimuli consisted of 16 directions (from 0° to 337.5°) and 2 spatial frequencies (0.03 and 0.13 cycles per degree) at 80% contrast. They were generated in PsychoPy 2.22. The full stimulus set was presented 5 times in one experiment in pseudo-random sequence. Gratings drifted at 1 Hz for 2 s, followed by a 4 s gray screen). Stimulus presentation onset was tracked using a photodiode and TTL pulses were generated and sampled by the imaging microscope to synchronize stimulation and imaging data.

### Analysis

#### Image processing

Scanbox .sbx files were converted to tiff format and motion corrected and segmented using Suite2p in Python (https://github.com/MouseLand/suite2p). The files from the contralateral and ipsilateral eye stimulation were aligned together to ensure the same cells were segmented. Files from different days were aligned and segmented separately. Rigid and non-rigid registration were run. After automated cell detection, the registered binary was manually checked and additional regions of interest were drawn if necessary.

#### Identification of visually responsive cells

The F for the entire fluorescent trace of each cell was calculated by subtracting 0.7* Fneuropil from Fcell. ΔF/F was calculated as (F-F0)/F0, where F0 was the baseline fluorescence. F0 was calculated as the 25^th^ percentile of the fluorescence signal in a 30 s sliding window (*84, 85*). ΔFstimulus/F was calculated as the ΔF/F during the stimulus window and ΔFoff/F was calculated as the ΔF/F during the gray screen presentation. To compute whether a neuron was significantly visually responsive, we performed a Wilcoxon Signed Ranks test comparing ΔFstimulus/F and ΔFoff/F with a Bonferroni corrected α = 0.05/ number of unique trials. Cells were considered visually responsive if they passed significance threshold for at least 25% of the trials of the single stimulus condition (*60, 86*). Cells were considered monocular if they had significant visual response to one or more stimulus conditions presented to either the contralateral or ipsilateral eye. Cells were considered binocular if they had a significant visual response to one or more stimulus conditions presented to each eye.

#### Ocular dominance index

The ocular dominance (ODI) of individual neurons was calculated as the ratio between the difference and the sum of the mean ΔFstimulus-peak/F in response to the ipsilateral or contralateral eye experimentally determined preferred drifting direction:

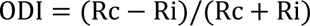

An ODI of -1 or +1 indicates ipsilateral or contralateral eye dominance, respectively. The stimulus-triggered response, ΔFstimulus-peak/F, was calculated by subtracting the mean ΔF/F for the 16 frames prior to stimulus presentation from the mean ΔF/F 7 to 31 frames (0.5 sec to 2 sec) after stimulus presentation (averaged 4 frames around the peak). Only neurons that were longitudinally tracked and responsive across imaging sessions were used for ODI calculations. For Fig. 4M, we used data only from longitudinally tracked neurons and binned the cells based on their pre MD ODI. We made 8 bins and calculated the median change in the binned cells’ contralateral eye responses and ipsilateral eye responses.

#### Direction and orientation tuning

The preferred orientation of a neuron was calculated as:

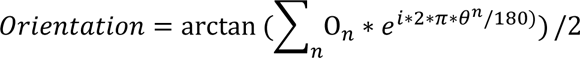

On is the peak neuronal response to the 18 different orientations (0° to 170° spaced every 18°).

Global orientation selectivity was calculated as:

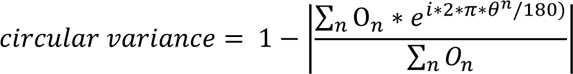

Cardinal proportions were calculated as the proportion of neurons having an orientation preference to 0° or 90°, ± 11.25°.

Binocular matching for binocular neurons was quantified as the absolute difference in preferred orientation of the contralateral and the ipsilateral eyes.

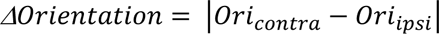

#### Spiking correlations

Using Suite2p, the deconvolved spike train was extracted from each cell. An unconstrained non-negative deconvolution using exponential kernels was used. The kernel decay timescale was set to 1.0 and a gaussian filter with a smoothing constant of 10 was applied to the neuropil subtracted fluorescence trace. The neuropil subtraction coefficient was set to 0.7. The spiking activity of each pair of significantly responsive, longitudinally tracked neurons was calculated by concatenating the spiking activity for all trial-on periods and performing a pairwise Pearson’s correlation. This was done independently for contralateral and ipsilateral eye responses.

#### Response reliability

For all responsive, longitudinally tracked neurons, the response reliability was calculated by the coefficient of variation (cv) of the neuron across all trials of each unique stimulus condition. *σ* is the standard deviation of the neuron’s responses to all trials of a specific stimulus and *μ* is the mean of the trial responses.

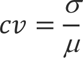

#### Longitudinal imaging

To locate the same imaging plane for longitudinal imaging, we used the images acquired on previous days as reference, such as the vascular map of the brain surface acquired with the PCO camera and epifluorescence and the mean motion-corrected two-photon fluorescence images. The angle of the objective was kept the same. Fine adjustment of imaging depth and x,y location was performed using a Scanbox built-in plugin ‘searchref’. Briefly, a zstack was automatically acquired and projected onto the mean motion-correct image from the prior imaging. The plugin computes (using fast fourier transform) the optimal translation of the microscope and moves the scope to best align the images (<u>Scanbox Searches for a Population – Scanbox</u>).

#### Longitudinal cell tracking

To track cells across multiple imaging sessions across different days, roiMatchPub.m, a MATLAB package written by Adam Ranson was used (https://github.com/ransona/ROIMatchPub). Briefly, the package uses a control point-based affine geometric transformation to correct for the plane rotation. The transformation is then applied to the image from the second imaging time point (“post MD”). The overlap of ROIs between the mask file from the reference image and the transformed mask file from the second imaging time point is calculated. Overlapping ROIs were considered as longitudinally tracked after visual inspection and verification (Extended Data Fig.9D and E). The average percent of cells that were segmented before MD that were longitudinally tracked and also segmented post MD was 46 ± 6% (Extended Data Fig.9F). For naïve mice, the average percent of cells that were longitudinally tracked from day 1 to day 6 of imaging was 26 ± 3.0% for WT mice and 35.9 ± 9.2% for CCN1 cKO mice (Extended Data Fig.11K). The difference in number of neurons that were longitudinally tracked between the two genotypes, combined with a lower number of mice in the WT naïve condition, could account for the difference in the unresponsive number of neurons for all segmented cells vs. longitudinally tracked unresponsive neurons (Extended Data Fig.11D and K).

#### Bootstrapping

For bootstrapping analysis in Fig. 4L, we performed a hierarchical bootstrapping method (*59*) with two levels, the first level being the animal and the second level the cells. Sampling was performed 10000 times for the dataset. A sample was taken with replacement at the first level (mice) with the sample size being equal to the number of mice. Then, a sample was taken with replacement at the second level, the neurons imaged. We chose a sample size of 300 which was slightly larger than the maximum number of neurons imaged per mouse. We reported the mean across the 10000 samples and directly calculated a p-value (pboot) that represents the probability that the mean ODI of the pre MD group is larger than that of the post MD group. The pboot was used to determine statistically significant differences for the pre MD and post MD samples. For the bootstrapping analysis in Fig. 4M, bottom panel, we performed a bootstrapping analysis with the bootci function in MATLAB for 1000 samples for each of the pre MD ODI bins. We calculated the mean for these 1000 samples and a 95% confidence interval.

### Statistical analysis

For most experiments, analyses were performed in GraphPad Prism. For WFA and PV quantifications and two-photon *in vivo* imaging data, analyses were run in MATLAB (MathWorks) and Python. Significance was set at alpha = 0.05 and corrected appropriately for multiple comparisons. Data were tested for normality, and two-tailed parametric or non-parametric tests were run as appropriate. Tests were corrected for multiple comparisons using either the Tukey method (for 2-way ANOVA). For non-parametric multiple comparisons, the Dunn method was applied. Adjusted p-values are shown. For chi-square tests of proportions done in MATLAB, a Bonferroni correction was applied for the number of tests. For chi-square tests, un-corrected p-values are shown, with significance indicated. For microglia morphology analyses, R studio was also used.

#### Data presentation

For the entirety of the study, bar charts show the average and error bars show the standard error of the mean (SEM). Box plots show the median and the upper and lower quartiles; the whiskers span the full data set. Violin plots show the median as the solid line and the upper and lower quartiles as dotted lines.

### Data availability

Developmental RNA sequencing is deposited at GEO GSE161398 and GSE99791. Other data available on reasonable request from the authors.

### Code availability

All code and analysis pipelines including the in vivo calcium imaging analysis, microglia morphology and classification, WFA quantification, and spiking correlations analysis is available at publication at: https://github.com/MNL-A/Sanchoetal2024.

## Acknowledgments

This work has been supported by NIH NINDS 1R01NS105742, L.I.F.E. Foundation, Chan Zuckerberg Initiative to N.J.A. L.S. was supported by NIH NEI NRSA 5F32EY033629 and the Salk Pioneer Fund. J.B. was supported by the NIH Blueprint and BRAIN Initiative D-SPAN Award 1F99NS134205. This work was supported by the Waitt Advanced Biophotonics Core Facility of the Salk Institute with funding from NIH-NCI CCSG: P30 CA01495, NIH-NlA San Diego Nathan Shock Center P30 AG068635, and the Waitt Foundation. This work was supported by the GT3 Core Facility of the Salk Institute with funding from NIH-NCI CCSG: P30 CA01495, an NINDS R24 Core Grant and funding from NEI (R24NS092943). We thank the members of the Allen laboratory, Dr. Edward Callaway, and Dr. Michael V. Sofroniew for providing feedback on the manuscript.

## Author Contributions

N.J.A. and L.S. conceived of the project and designed the experiments. L.S. performed the experiments, designed the analysis pipelines, and conducted the data analysis. M.M.B. performed the RNA sequencing experiments. J.B. performed the analysis and generated plots for the RNA sequencing. T.D. and E.W. contributed to immunostaining and image analysis. L.S. and N.J.A. interpreted the results and wrote the paper with comments from all authors.

## Materials & Correspondence

Correspondence and request for materials should be addressed to Nicola J. Allen.

## Supplementary Data Tables

**Supplemental Data Table 1.** Data from the RNA sequencing experiments. Developmental data from Farhy-Tselnicker et al. 2021 and Boisvert et al. 2018. *Ccn1* is named *Cyr61*.

**Supplemental Data Table 2.** Gene-set enrichment analysis (GSEA) data. Enrichment of gene sets and pathways from the Gene Ontology (GO), Reactome, KEGG and WikiPathway. Only genes that had an FPKM > 1 in each sample were included. Genes were ranked based on descending log2 Fold Change (LFC) in each comparison. A cutoff of adjusted p value < 0.05 was used to determine significantly enriched pathways and terms.

**Supplemental Data Table 3.** Overrepresentation analysis (ORA) showing GO terms that were enriched in the significant DEGs across the plasticity paradigms. Criteria for selection of genes: FPKM > 1 in each sample of at least one group per comparison. The following criteria were used to select significantly enriched GO terms: adjusted p value < 0.05 (Benjamini and Hochberg correction) and q-value < 0.01.

**Supplemental Data Table 4.** Statistical tests and summary statistics from studies.

---

**Extended Data Fig. 1.**
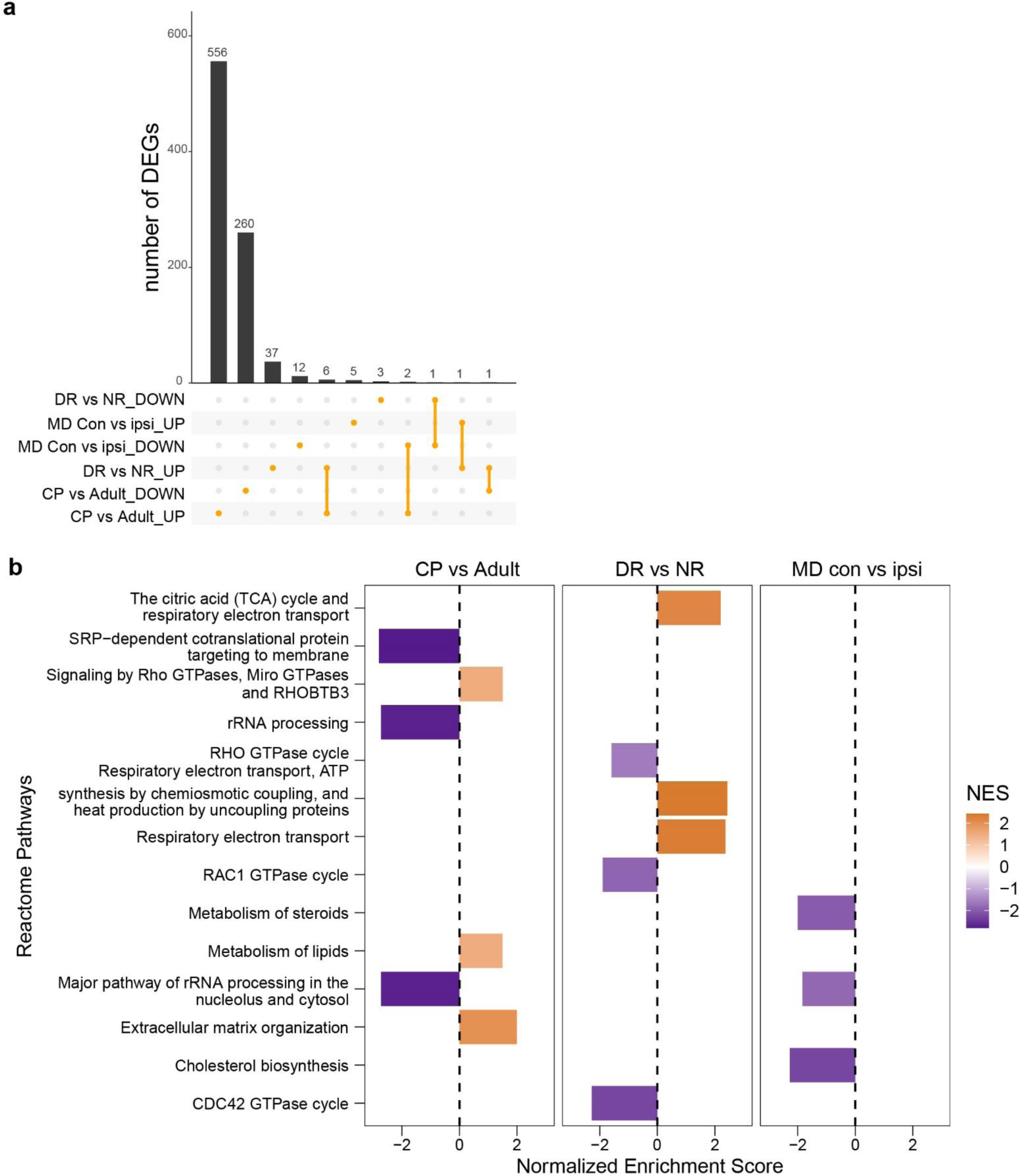
Transcriptomic analysis of visual cortex astrocytes across development and plasticity. Upset plot showing up- and down-regulated genes across the different comparisons (MD con vs ipsi, DR vs NR, CP vs Adult). MD, monocular deprivation. Con, contralateral eye to deprivation. Ipsi, ipsilateral eye to deprivation. CP, critical period. Adult, P120. DR, dark rearing. NR, normal rearing. (criteria: FPKM > 1 in at least one group per comparison, ribotag pulldown FPKM (astrocyte) /input FPKM (all cells) > 0.75 in at least one group per comparison, adjusted p-value < 0.05, Log2 Fold Change (LFC) > |0|). **b**. Gene-set enrichment analysis (GSEA) results using the Reactome Database (ranked gene lists based on LFC only. Significant (adjusted p-value < 0.05) Reactome Pathways were first selected if they were found in at least two comparisons. The Pathways were then ranked by Normalized Enrichment Score (NES) and the top 6 (three positive and three negative enrichment scores) were plotted for each comparison.

**Extended Data Figure 2.**
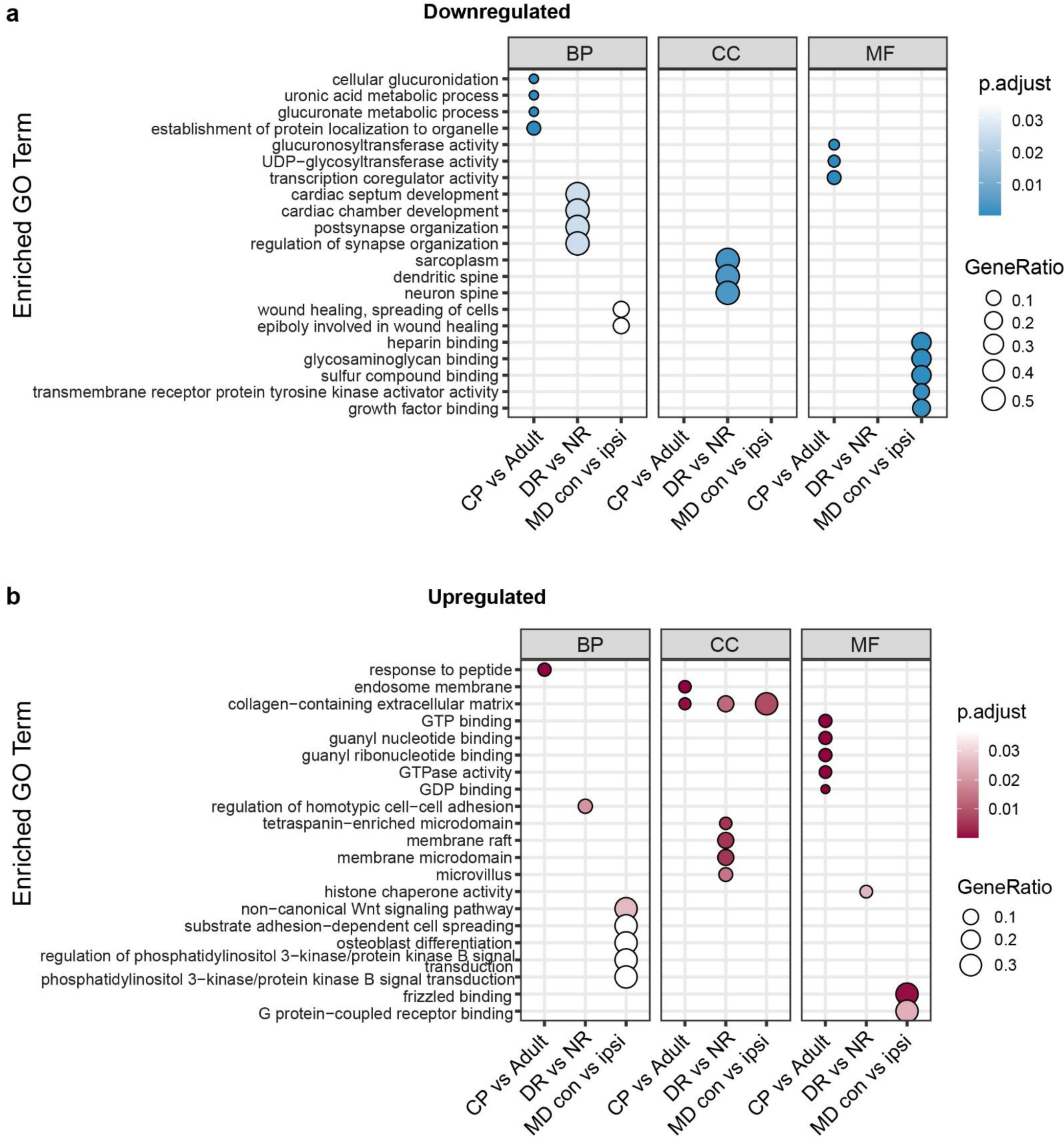
Overrepresentation analysis (ORA) to determine Gene Ontology (GO) terms enriched in DEGs across the comparisons. **a**. Enriched GO terms for downregulated genes. **b**. Enriched GO terms for upregulated genes. Size of circles denotes the Gene Ratio (number of DEGs from each gene list found in the GO Term). Color shade denotes adjusted p value. BP: Biological Process, CC: Cellular Component, MF: Molecular Function.

**Extended Data Figure 3.**
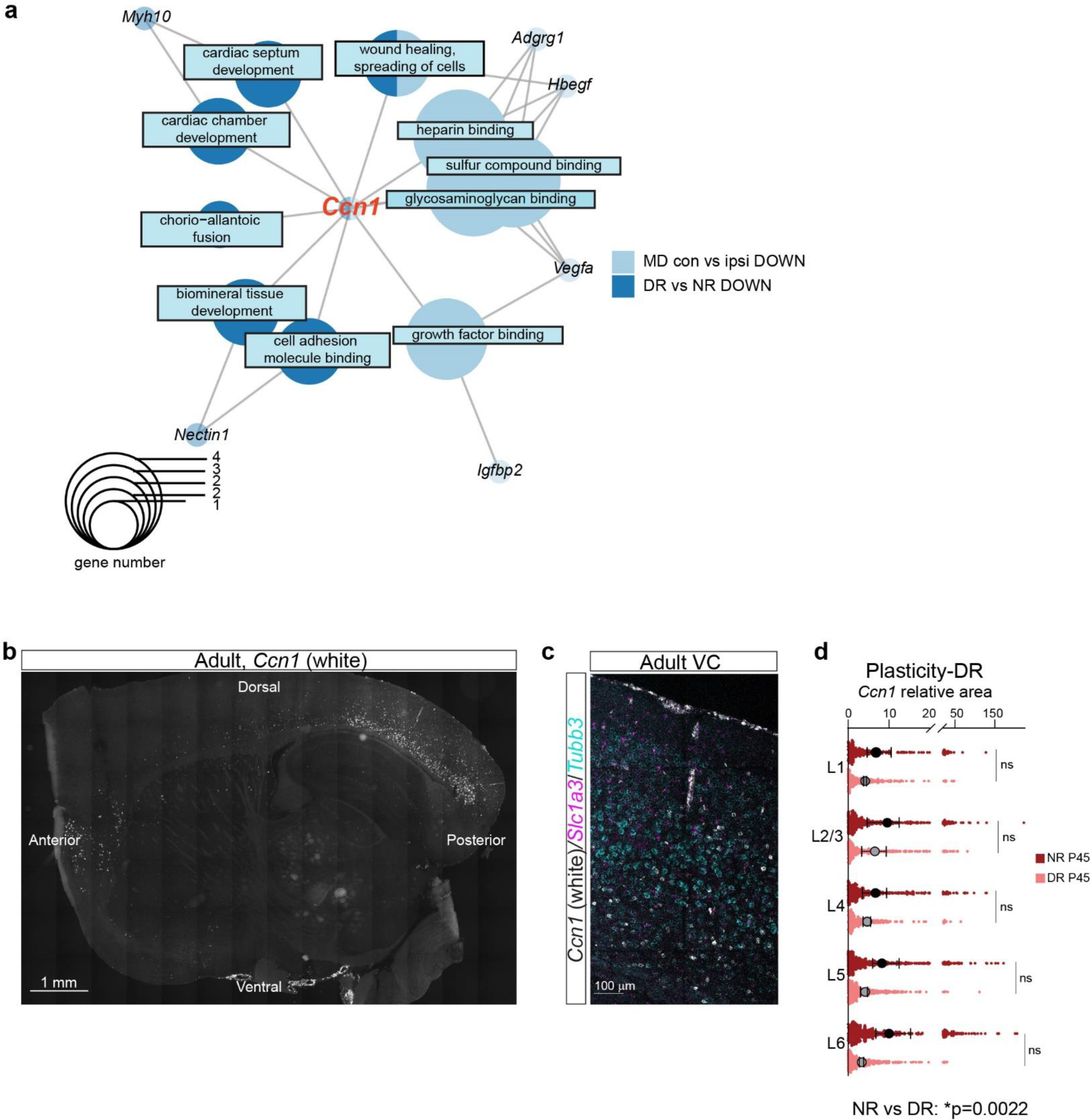
Identification of *Ccn1* as a candidate factor and validation of RNAseq using smFISH. **a**. *Ccn1* overrepresentation analysis network example. Light/dark blue circles with text boxes represent ORA networks downregulated in MD con vs ipsi and DR vs NR, respectively. Size of the circle denotes number of genes. **b**. Tiled sagittal section of an adult mouse smFISH for *Ccn1*. Scale bar, 1 mm. **c.** Representative image of P120 smFISH in a tile image of the visual cortex (VC). smFISH against *Slc1a3, Tubb3,* and *Ccn1.* Same image from Fig.1H. **d**. *Ccn1* relative area per astrocyte after dark-rearing to P45 (DR) or normal rearing to P45 (NR) in the VC. Two-way ANOVA. Small dots denote individual astrocytes, large circles are averages by mouse. N = 3 mice.

**Extended Data Figure 4.**
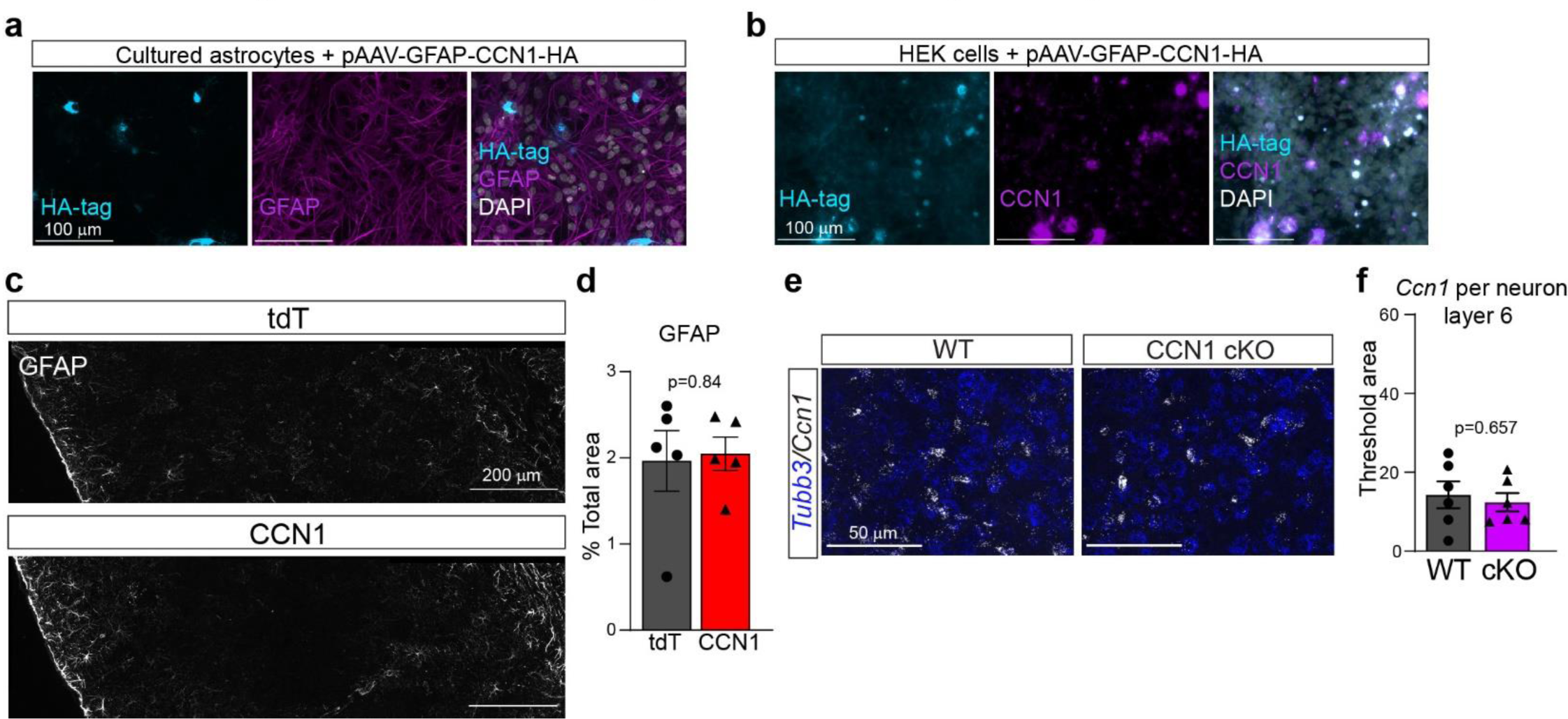
Validation of *in vivo* approaches to astrocytic manipulation of CCN1. **a**. Immunocytochemistry (ICC) of astrocytes transfected with pAAV-GFAP-CCN1-HA. Cyan, HA-tag. Scale bar, 100 µm. **b**. ICC of HEK cells transfected with pAAV-GFAP-CCN1-HA. Scale bar, 100 µm. **c**. Representative image of visual cortex (VC) GFAP immunostaining in tdT and CCN1 infected P28 mice. Scale bar, 200 µm. **d**. GFAP quantified as % total area of section. N = 5 mice in tdT, 5 mice in CCN1. Unpaired T-test **e**. Representative image of *Ccn1* smFISH in layer 6 VC of adult WT and CCN1 cKO mice. Scale bar, 50 µm. **f**. *Ccn1* threshold area per *Tubb3*+ cell in layer 6 of VC. Unpaired T-test. For **d** and **f,** black symbols denote mouse averages.

**Extended Data Figure 5.**
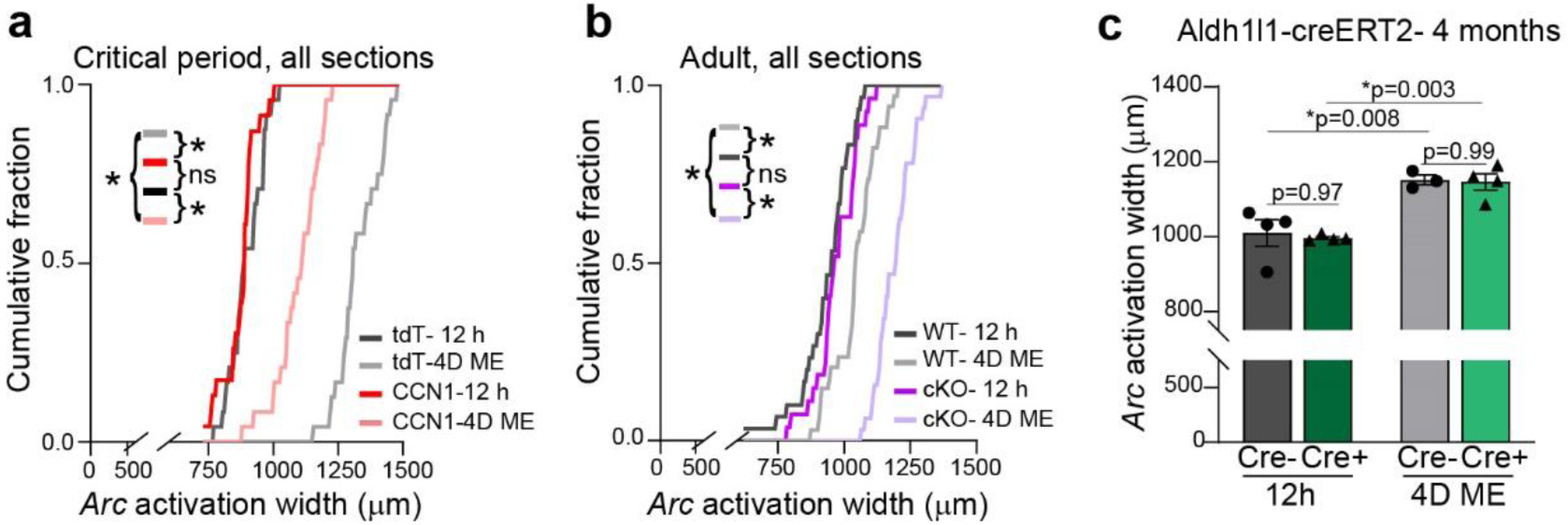
Astrocyte CCN1 regulates binocular zone remodeling. **a**. Cumulative probability distribution of all sections *Arc* induction assay in critical period mice after 12 hours of monocular enucleation (12 h) or 4 days of monocular enucleation (4D ME). 23-24 sections per group. Kolmogorov-Smirnov tests. **b**. As in **a** but in 4 month WT and cKO mice. 27-34 sections per group. Kolmogorov-Smirnov tests. **c**. *Arc* induction assay in 4 month old Aldh1l1-creERT2 mice, not crossed to *Ccn1*^fl/fl^ mice. Two-way ANOVA with post-hoc Tukey tests. Black symbols denote mouse averages. Cre-12 h, 4 mice. Cre+ 12 h, 4 mice. Cre-4D ME, 3 mice. Cre+ 4D ME, 4 mice.

**Extended Data Fig.6.**
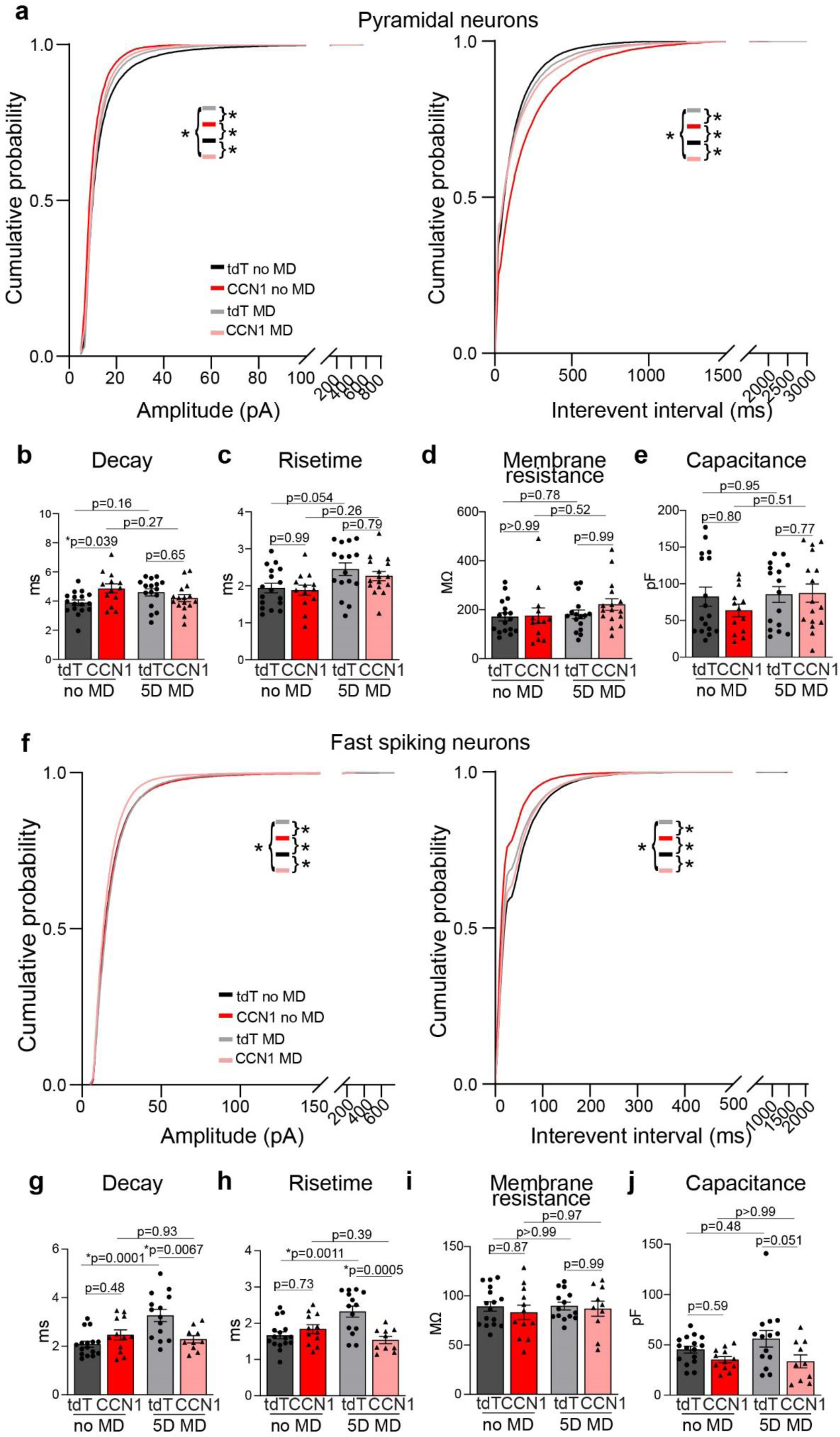
Astrocyte CCN1 regulates excitation on layer 2/3 visual cortex neurons. **a**. Cumulative probability distribution of amplitude (left) and interevent interval (right) for all events in pyramidal neurons. Kolmogorov-Smirnov (KS) tests. **b**. Average decay. **c**. Average 10-90% risetime. **d**. Membrane resistance (MΩ). **e**. Capacitance (pF). tdT no MD: n = 17 cells, 9 mice. tdT MD: n = 16 cells, 7 mice. CCN1 no MD: n = 13 cells, 7 mice. CCN1 MD: n = 16 cells, 6 mice. **f**. Cumulative probability distributions of amplitude (left) and interevent interval (right) for all events in fast-spiking neurons. KS tests. **g**. Average decay. **h**. Average 10-90% risetime. **i**. Membrane resistance. **j**. Capacitance. All statistics, excluding KS tests are two-way ANOVAs with post-hoc Tukey tests. Black symbols denote individual cells.

**Extended Data Figure 7.**
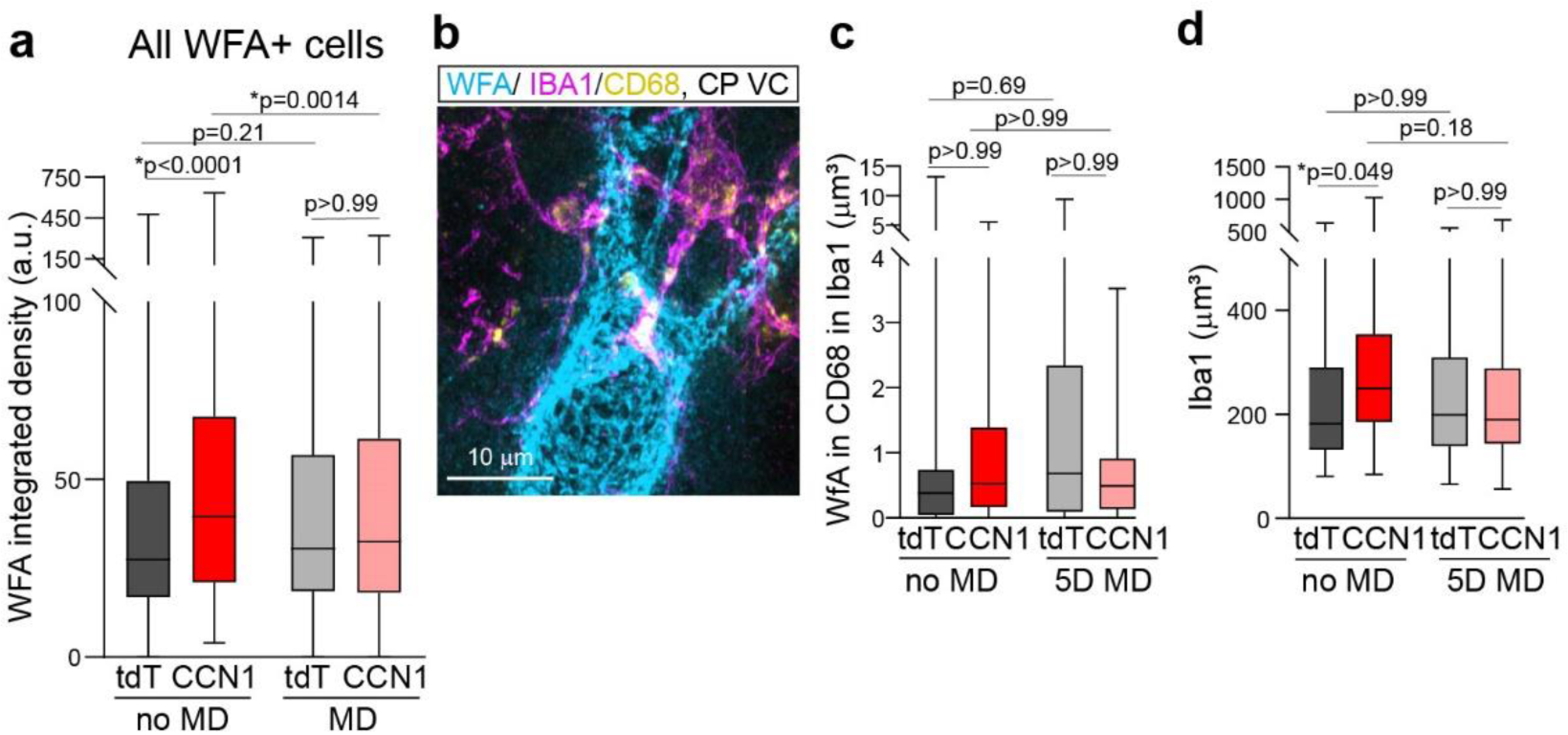
Astrocytic CCN1 regulates PNN density and microglial activation during the critical period. **a**. WFA integrated density in all WFA+ cells, not just PV cells. tdT no MD = 577 cells. tdT MD = 572 cells. CCN1 no MD = 491 cells. CCN1 MD = 538 cells. **b**. Representative image of IBA1, WFA, and CD68 staining in critical period VC. Scale bar, 10 µm. **c**. Volume of WFA within CD68+ puncta within IBA1 volume. **d**. IBA1 volume with field of view (FOV). N = 39-40 FOVs per group from 4 mice per group. All statistics are Kruskal Wallis with post-hoc Dunn’s tests shown on graphs.

**Extended Data Fig. 8.**
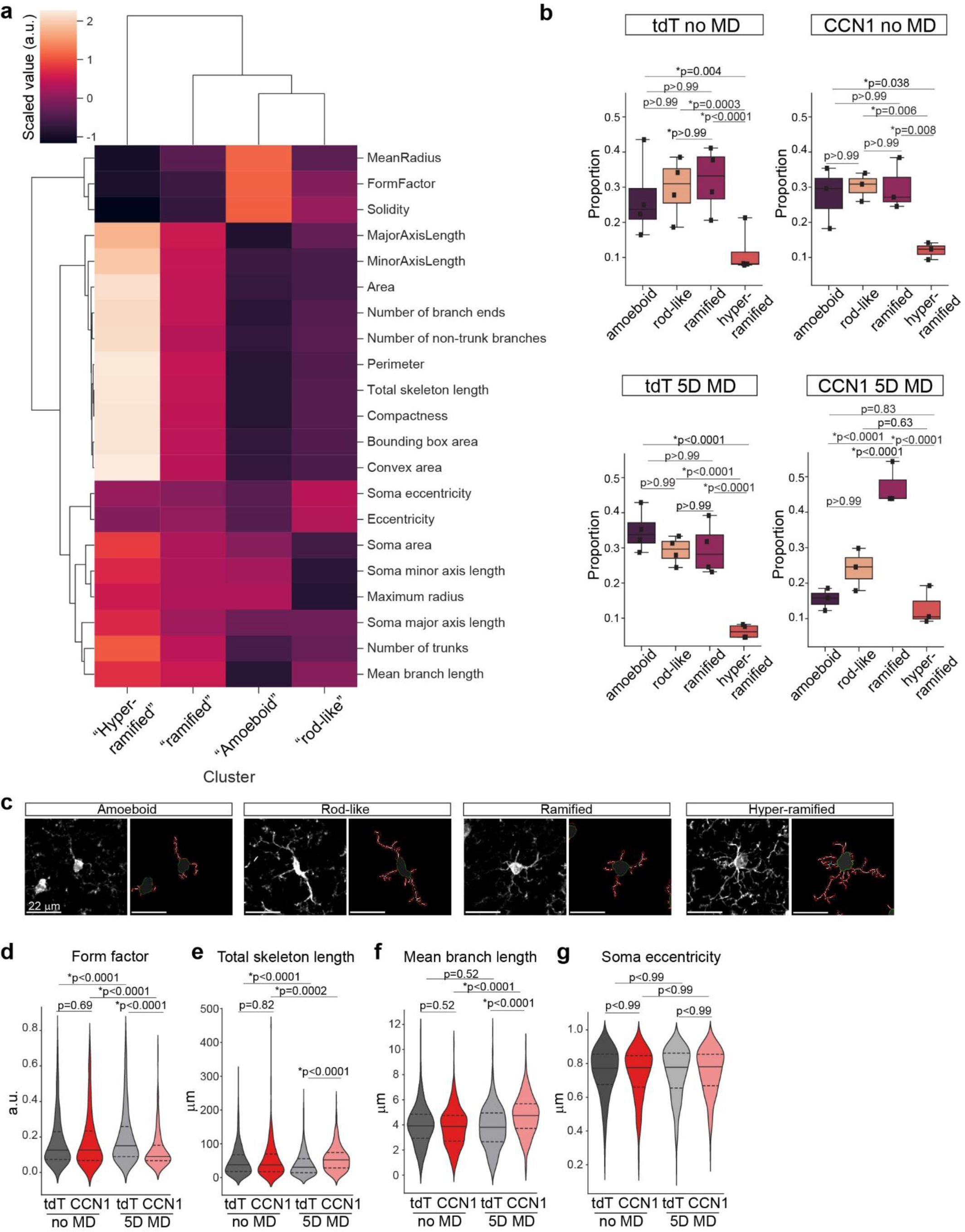
Astrocytic CCN1 regulates microglial morphology during the critical period. Microglial morphology in critical period age mice. **a**. Clustered heatmap of standard scaled features for each defined microglial morphotype. Color bar is scaled values, arbitrary units (au). **b**. Proportion of microglial morphotypes-amoeboid, rod-like, ramified, and hyper-ramified separated by experimental group. N = 3-4 mice per group. Black symbols, mouse averages. 2-way ANOVA with post-hoc Tukey’s tests. **c**. Representative images and skeletons of amoeboid microglia, rod-like, ramified, and hyper-ramified, left to right. **d**. Form factor (4*π*Area/Perimeter^2^), a measure of circularity. **e**. Total skeleton length. **f**. Mean branch length. **g**. Soma eccentricity. **c-f**. tdT no MD, 747 microglia from 4 mice. tdT MD, 876 microglia from 4 mice. CCN1 no MD, 496 microglia from 3 mice. CCN1 MD, 379 microglia from 3 mice. All statistics are Kruskal Wallis with post-hoc Dunn’s tests shown on graphs.

**Extended Data Fig. 9.**
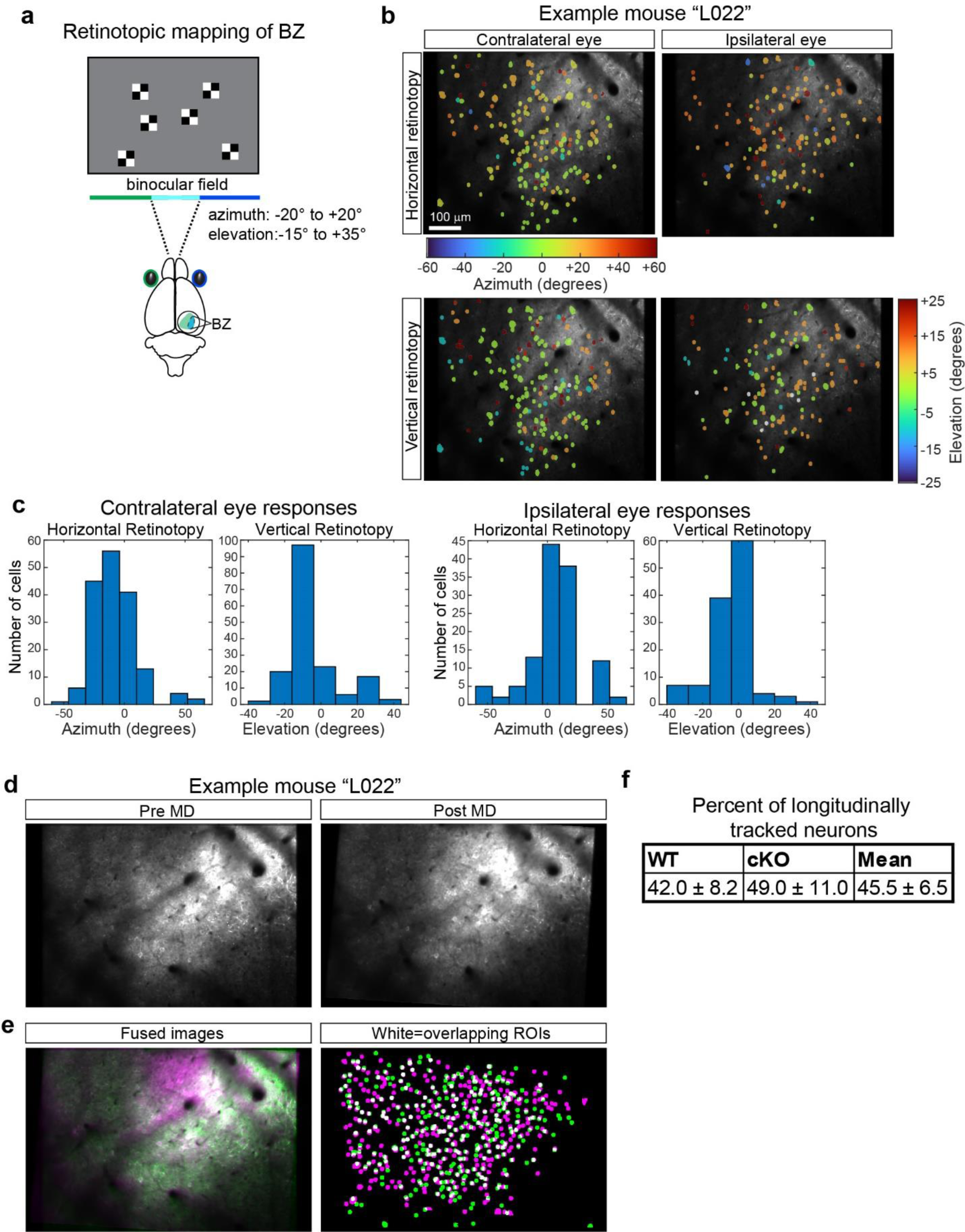
Validation of 2-photon approach. Validation of 2-photon approach. **a**. Prior to ODI mapping, retinotopic mapping of the BZ is performed with small checkerboard boxes. Stimuli are presented independently to either eye. **b**. Representative retinotopy in degrees of visual field in example mouse “L022”. Top, horizontal retinotopy, “azimuth”. Bottom, vertical retinotopy, “elevation”. Left, contralateral eye responses. Right, ipsilateral eye responses. Scale bar, 100 µm. **c**. Example mouse “L022”. Histograms of number of cells with peak responses to a particular azimuth (horizontal) or elevation (vertical). Regions are considered binocular if peak responses are within -20° to +20 ° in azimuth and -15° to +35 ° in elevation for both eyes. **d**. Validation of longitudinal tracking using roiMatchPub pipeline in MATLAB. Example mouse “L022”. Mean images pre, left, and post, right, MD. **e**. Left, fused image. Right, white denote overlapping ROIs. **f**. Percent of longitudinally tracked cells across WT and CCN1 cKO mice.

**Extended Data Fig. 10.**
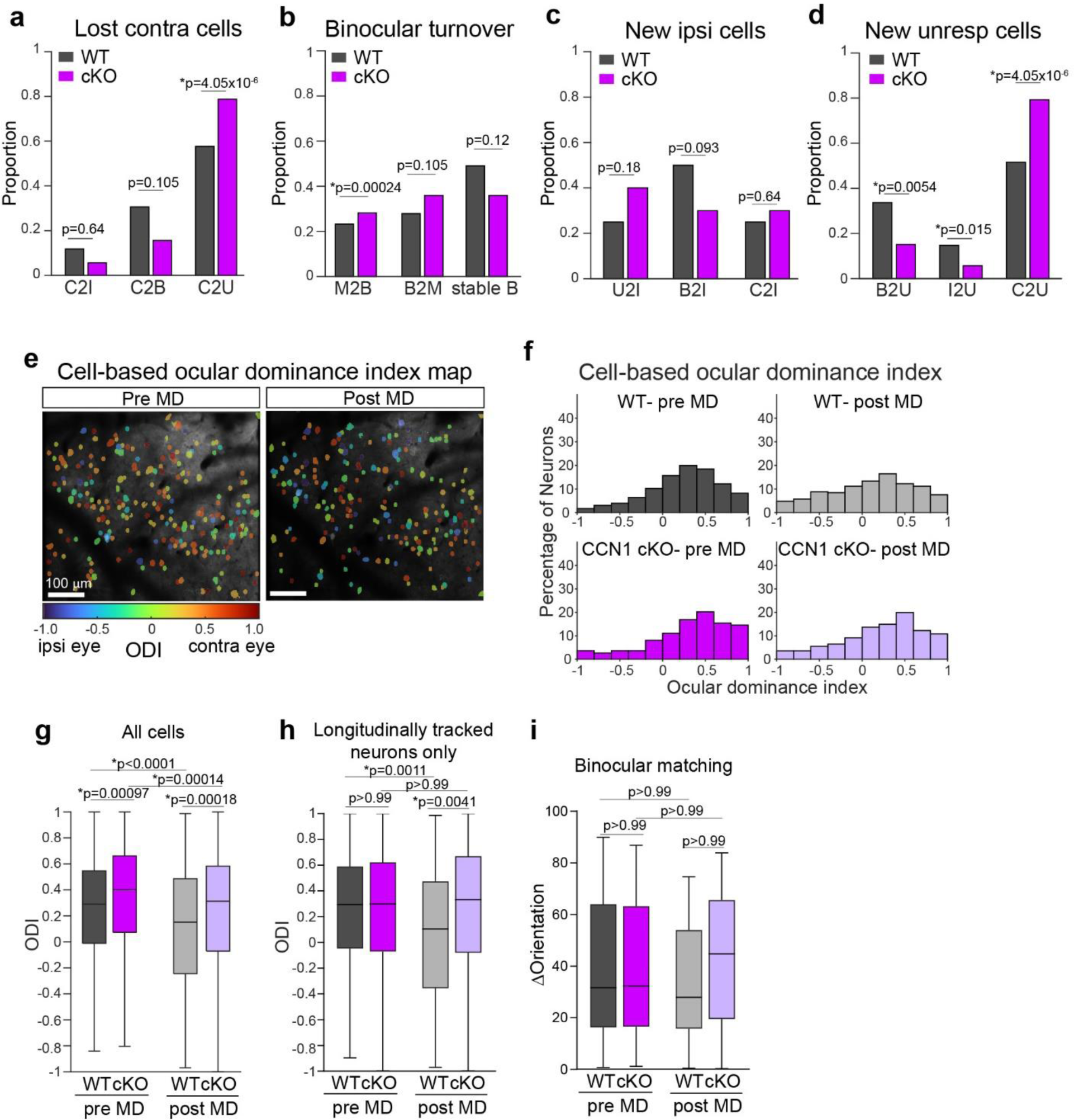
Astrocyte CCN1 cKO mice have aberrant binocular circuitry. Astrocyte CCN1 cKO mice have aberrant binocular circuitry. **a**. Contralateral cell loss. C2I, contralateral to ipsilateral. C2B, contralateral to binocular. C2U, contralateral to unresponsive. **b**. Binocular turnover. B2M, binocular to monocular. M2B, monocular to binocular. Stable B, stable binocular. **c**. New ipsilateral cells. C2I, contralateral to ipsilateral. B2I, binocular to ipsilateral. U2I, unresponsive to ipsilateral. Proportions are expressed as a fraction of newly formed ipsilateral cells after MD. **d**. New unresponsive cells. I2U, ipsilateral to unresponsive. C2U, contralateral to unresponsive. B2U, binocular to unresponsive. Proportions are expressed as a fraction of newly formed unresponsive cells after MD. **a-d**. Chi-square tests. Correction for multiple comparisons (adjusted p-value threshold = 0.01667). Stars denote significance. **e**. Cell-based ocular dominance index map. Representative image with ocular dominance index color coded. Cool colors denote ipsilateral eye preference, warm colors denote contralateral eye preference. Scale bar, 100 µm. **f**. Histograms of ocular dominance index of all cells as a percentage of responsive neurons, regardless of longitudinal tracking. **g**. Box plots of data in **f**. WT-pre MD = 682 cells. WT-post MD = 554 cells. cKO-pre MD = 569 cells. cKO-post MD = 736 cells. **h**. ODI of longitudinally imaged neurons only. WT = 223 cells. cKO = 163 cells. **i.** Binocular matching as difference in preferred orientation for contralateral and ipsilateral responses for longitudinally tracked binocular neurons. WT = 72 cells. cKO =37 cells. Data from 4 mice per group. **g-i**. Kruskal-Wallis tests with post-hoc Dunn’s tests.

**Extended Data Fig. 11.**
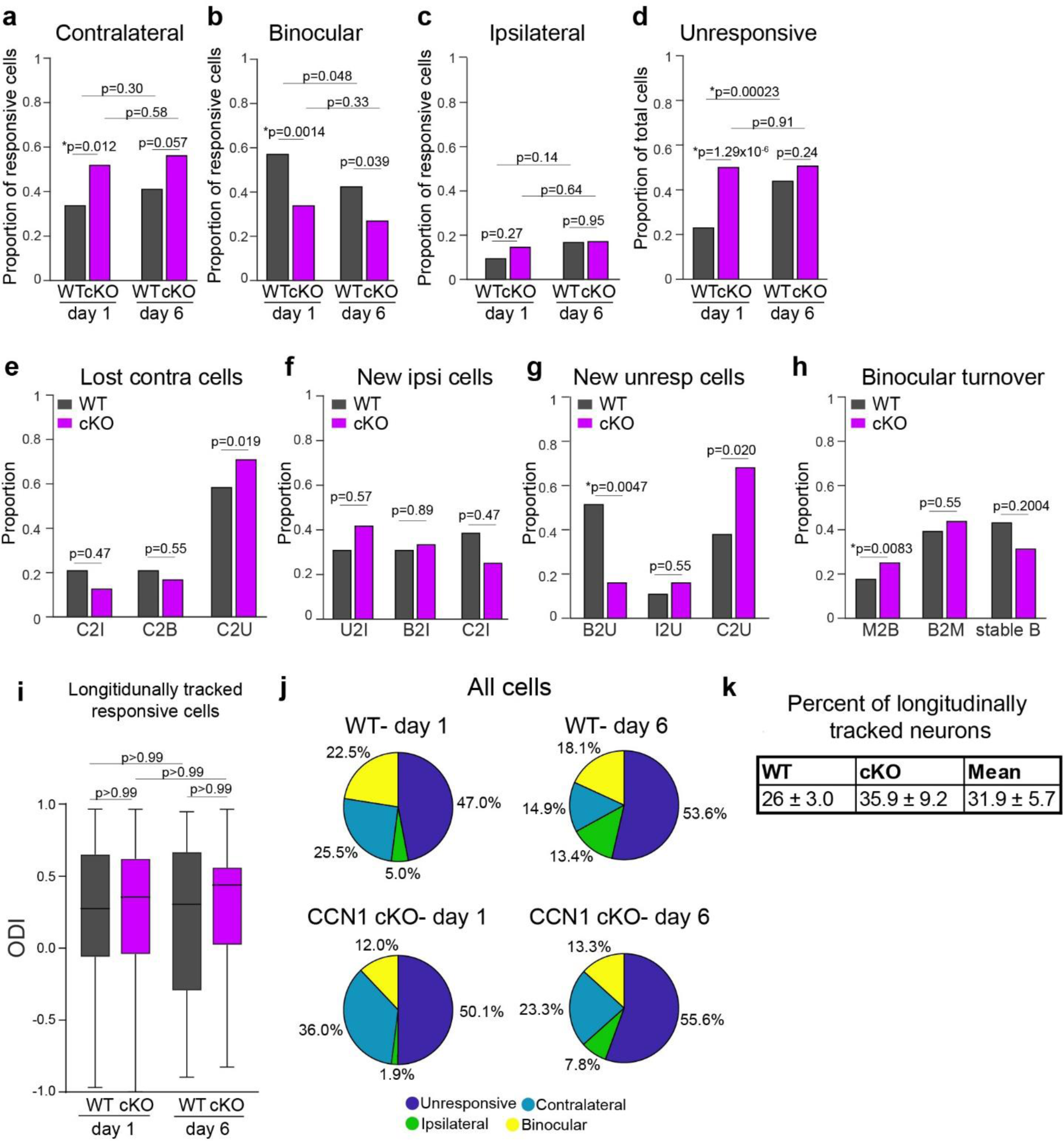
Naive CCN1 cKO mice demonstrate increased binocular cell turnover. Naïve CCN1 cKO mice demonstrate circuit instability. a-i. Longitudinally tracked neurons. a. Contralateral cells as a percentage of responsive cells. b. Binocular cells as a percentage of responsive cells. c. Ipsilateral cells as a percentage of responsive cells. a-c. WT responsive cells day 1 = 107. WT responsive cells day 6 = 78. CCN1 cKO responsive cells day 1 = 83. CCN1 cKO responsive cells day 6 = 82. **d**. Unresponsive cells as a proportion of total cells. WT total cells = 139. CCN1 cKO total cells = 166. **e**. Contralateral cell loss. C2I, contralateral to ipsilateral. C2B, contralateral to binocular. C2U, contralateral to unresponsive. **f**. New ipsilateral cells. C2I, contralateral to ipsilateral. B2I, binocular to ipsilateral. U2I, unresponsive to ipsilateral. Proportions expressed as a ratio of newly formed ipsilateral cells at day 6 of imaging. **g**. New unresponsive cells. I2U, ipsilateral to unresponsive. C2U, contralateral to unresponsive. B2U, binocular to unresponsive. Proportions expressed as a ratio of newly formed unresponsive cells at day 6 of imaging. **h**. Binocular turnover. B2M, binocular to monocular. M2B, monocular to binocular. Stable B, stable binocular. **a-h**. Chi-square tests of proportions. Stars denote significance. Correction for multiple comparisons (**a-d**, adjusted p-value threshold = 0.0125. **e-h**, adjusted p-value threshold = 0.01667). **i**. Ocular dominance index for longitudinally tracked responsive cells. WT, 64 cells. cKO, 52 cells. **j**. Pie chart of cell identity proportions for all segmented neurons, regardless of whether they were longitudinally tracked or not. **k**. Percent of longitudinally tracked neurons in WT and cKO naïve mice. Data from 2 WT mice and 3 CCN1 cKO mice.

**Extended Data Fig. 12.**
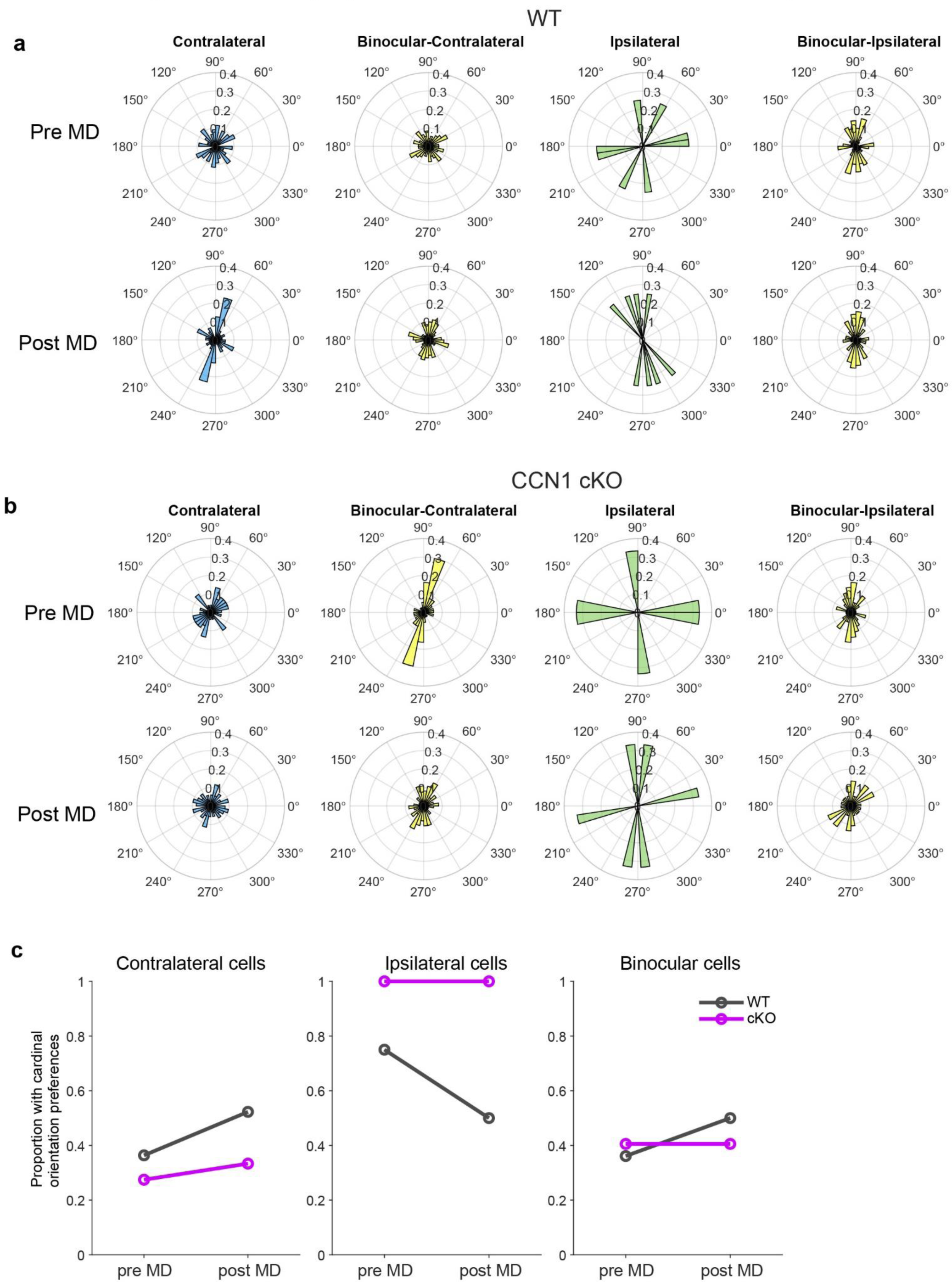
Cardinal proportions are not different in WT or astrocyte CCN1 cKO mice. Cardinal orientation proportions for longitudinally tracked contralateral, binocular, and ipsilateral neurons. **a**. Polar histograms for WT mice of preferred orientations for contralateral eye responsive neurons (blue), binocular neurons-contralateral responses (yellow), ipsilateral eye responsive neurons (green), and binocular neurons-ipsilateral responses. Top, Pre MD. Bottom, Post MD. Histograms are mirrored from 180° to 360°. Contralateral neurons = 56. Binocular neurons = 72. Ipsilateral neurons = 4. **b**. As in (A) but for CCN1 cKO mice. Contralateral neurons = 51. Binocular neurons = 37. Ipsilateral neurons = 3. **c**. Relative proportions of contralateral, ipsilateral, and binocular neurons with orientation preferences that are one of the cardinal orientations (0°, 90°, 180°). Chi-square tests. Contralateral neurons - WT pre vs post, p = 0.43; cKO pre vs post, p = 0.52; WT pre vs cKO pre, p = 0.47; WT post vs cKO post, p = 0.41. Ipsilateral neurons – WT pre vs post, p = 0.47; cKO pre vs post, not enough degrees of freedom; WT pre vs cKO pre, p = 0.35; WT post vs cKO post, p = 0.15. Binocular neurons – WT pre vs post, p = 0.092; cKO pre vs post, p > 0.99; WT pre vs cKO pre, p = 0.59; WT post vs cKO post, p = 0.35. For binocular neurons, contralateral and ipsilateral response cardinal proportions were averaged.

**Extended Data Figure 13.**
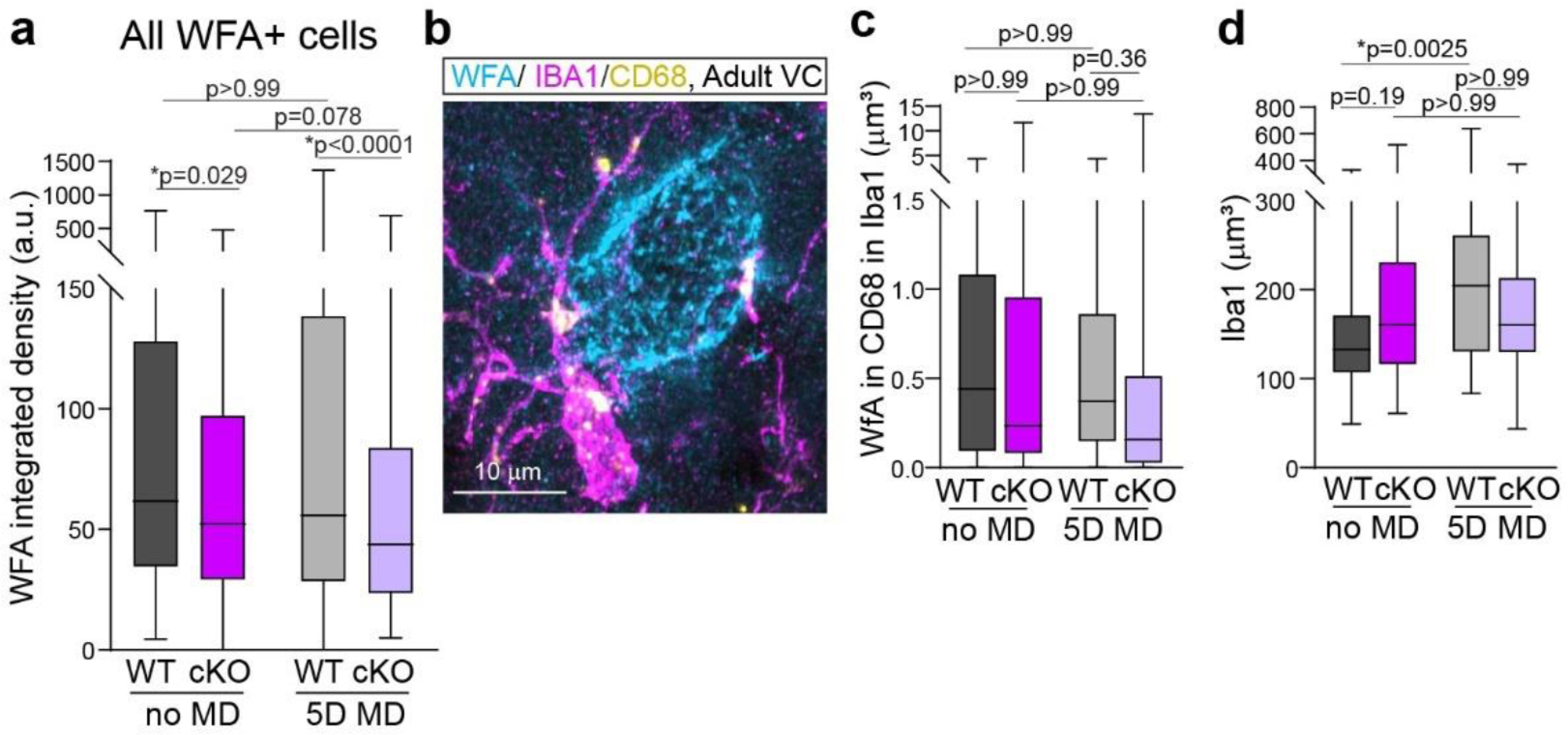
WFA and PNNs in adult mice. **a**. WFA integrated density quantified per WFA+ cell. WT no MD-493 cells. WT MD-506 cells. cKO no MD-474 cells. cKO MD-446 cells. **b**. Representative image of IBA1, WFA, and CD68 staining in adult VC. Scale bar, 10 µm. **c**. WFA colocalized to CD68+ puncta within IBA1 volumes. **d**. IBA1 volume within each field of view (FOV). N=39-40 FOVs per group from 4 mice per group. All statistics are Kruskal Wallis with posthoc Dunn’s tests shown on graphs.

**Extended Data Figure 14.**
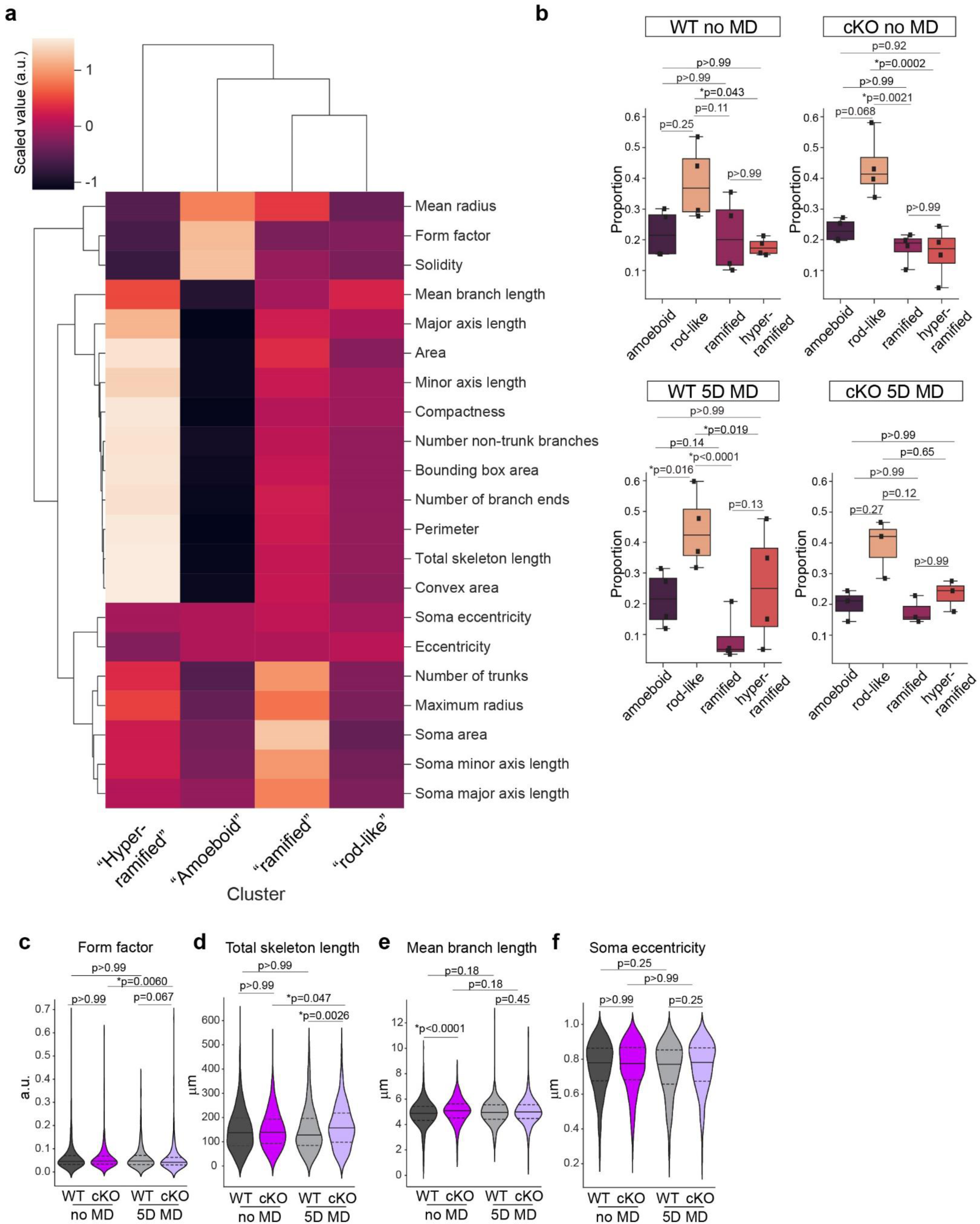
Microglial morphology in adults. **a**. Clustered heatmap of scaled features for each defined microglial morphotype. **b**. Proportion of microglial morphotypes-amoeboid, rod-like, ramified, and hyper-ramified separated by experimental group. N = 3-4 mice per group. Black symbols, mouse averages. 2-way ANOVA with post-hoc Tukey’s tests. **c**. Form factor. **d**. Total skeleton length. **e**. Mean branch length. **f**. Soma eccentricity. **c-f**. WT no MD, 883 microglia from 4 mice. WT MD, 612 microglia from 4 mice. cKO no MD, 667 microglia from 4 mice. cKO MD, 543 microglia from 3 mice. All statistics are Kruskal Wallis with posthoc Dunn’s tests shown on graphs.

